# Designing of epitope-based vaccine from the conserved region of spike glycoprotein of SARS-CoV-2

**DOI:** 10.1101/2020.08.27.269456

**Authors:** Vidhu Agarwal, Pritish Varadwaj, Akhilesh Tiwari

## Abstract

The emergence of COVID-19 as a pandemic with a high morbidity rate is posing serious global concern. There is an urgent need to design a suitable therapy or vaccine that could fight against SARS-CoV-2 infection. As spike glycoprotein of SARS-CoV-2 plays a crucial role in receptor binding and membrane fusion inside the host, it could be a suitable target for designing of an epitope-based vaccine. SARS-CoV-2 is an RNA virus and thus has a property to mutate. So, a conserved peptide region of spike glycoprotein was used for predicting suitable B cell and T cell epitopes. 4 T cell epitopes were selected based on stability, antigenicity, allergenicity and toxicity. Further, MHC-I were found from the immune database that could best interact with the selected epitopes. Population coverage analysis was also done to check the presence of identified MHC-I, in the human population of the affected countries. The T cell epitope that binds with the respective MHC-I with highest affinity was chosen. Molecular dynamic simulation results show that the epitope is well selected. This is an in-silico based study that predicts a novel T cell epitope from the conserved spike glycoprotein that could act as a target for designing of the epitope-based vaccine. Further, B cell epitopes have also been found but the main work focuses on T cell epitope as the immunity generated by it is long lasting as compared to B cell epitope.

## Introduction

Coronavirus infection is initiated with the interaction between the viral envelope and the host cellular membrane^1^. The viral envelope is made up of spike (S) and membrane (M) glycoproteins as well as envelope (E) non-glycosylated protein. S protein is a very important protein that is responsible for receptor binding, membrane fusion, internalization of the virus, tissue tropism and host range. Thus, S protein can be an appropriate target for vaccine development^2^. As there is an urgent need to fight with SARS coronavirus 2 (SARS-CoV-2) infection and vaccine are a cost-effective way to control it^3^.

## 2. MATERIALS AND METHOD

The following flowchart describes the method as seen in Figure 1.

**Figure1:**
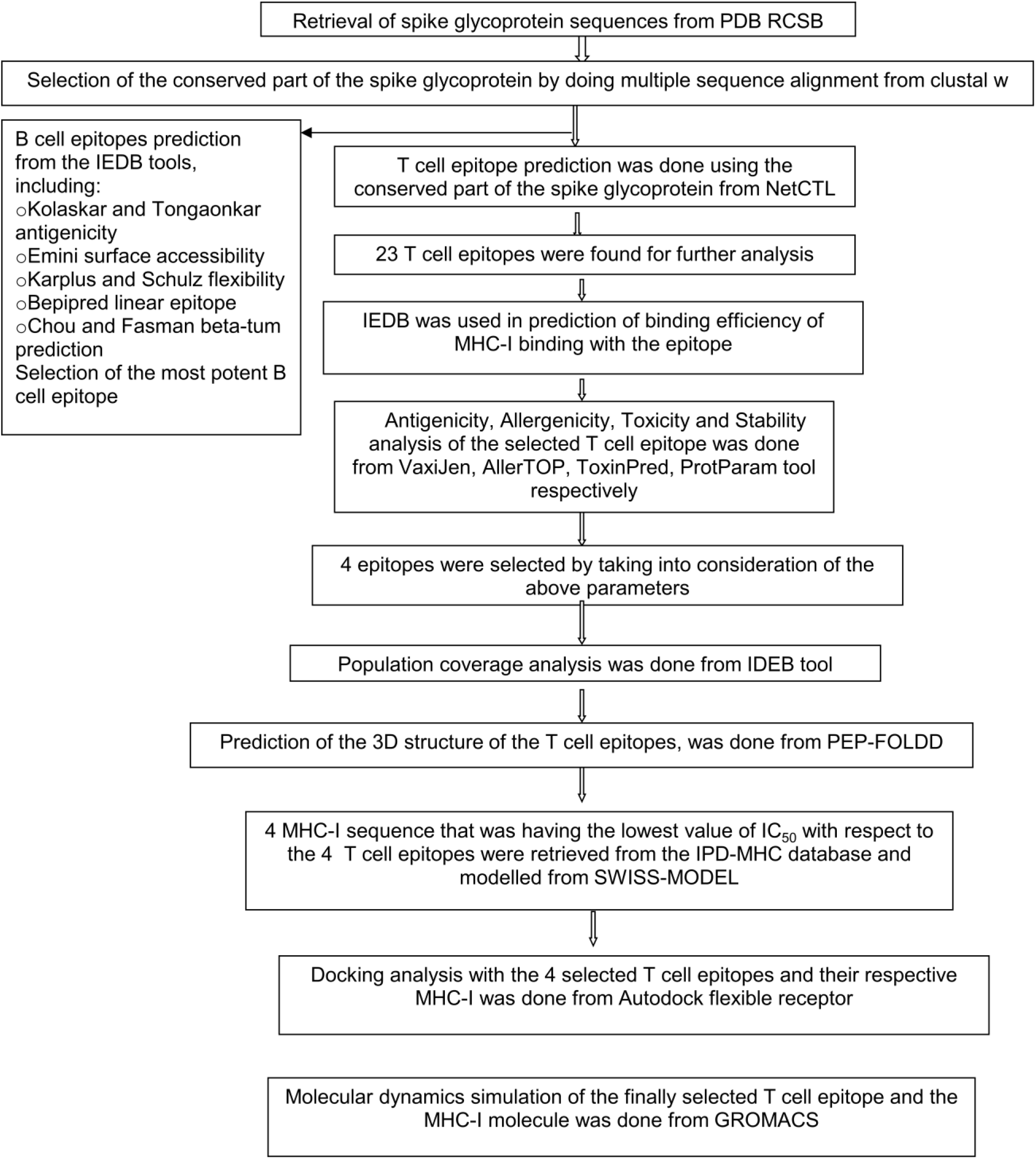
Shows the flowchart of the work done.

### 2.1. S glycoprotein sequence retrieval

S glycoprotein (outer membrane) sequences of SARS-CoV-2 were retrieved from Regional collaboratory for structural bioinformatics protein data bank (RCSB PDB) database, in FASTA format for further analysis.

### 2.2. Multiple sequence alignment (MSA)

MSA predicted the conserved region between different S glycoprotein sequences of SARS-CoV-2 that were available on PDB RCSB databank, using ClustalW.

### 2.3. T-cell epitope prediction

NetCTL1.2 found T-cell epitopes from the conserved S glycoprotein sequence of SARS-CoV-2, which have the potency to elicit an immune response. This prediction method included Major histocompatibility complex-I (MHC-I) binding along with TAP transport efficiency and proteasomal C-terminal (CT) cleavage. This type of epitope prediction is done taking into consideration of 12 MHC supertypes. MHC class I binding and proteasomal CT cleavage is done by using artificial neural network (ANN) method, whereas TAP transport efficiency is detected by using weight matrix.

The parameters used in the analysis were set as follows: Threshold of 0.5 to maintain the sensitivity of 0.89 and specificity of 0.94, Supertype A1, Weight on CT and TAP transport efficiencies of 0.15 and 0.05, respectively.

Further, the Immune epitope database (IEDB) tool predicted MHC-I binding with half maximal inhibitory concentration (IC_50_) values of the T-cell epitopes. MHC-I molecule binding, was calculated using Stabilized matrix-based method (SMM). Allele selection was done and the lengths of the epitopes were set to be 9.0, for proceeding with the binding analysis. This tool finally produced scores for the proteosomal processing, MHC-I binding, TAP transport and an overall score, which indicates the peptides intrinsic potential of a T-cell epitopes^4^.

### 2.4. Physiochemical analysis of T cell epitopes

T cell epitope prediction that are effective in eliciting an immune response and are safe for the host, can save a lot of wet lab efforts. The following are the physiochemical properties that must be present for an effective, safe and stable T cell epitope: Antigenicity, Non-allergenicity, Non-toxicity and stability. Based on these properties, 4 T cell epitopes were selected from the 23 T cell epitopes selected from the immune database.

#### 2.4.1. Antigenicity determination of T cell epitopes

Antigenicity determination of T cell epitopes means that whether the epitope is capable of eliciting an immune response or not, inside the host. VaxiJen v2.0 predicts the protective antigens and vaccine subunits

#### 2.4.2. Allergenicity determination of T cell epitopes

Allergenicity determination of the epitopes check whether the epitope is producing any kind of allergic reactions or hypersensitivity or not. AllerTOP v. 2.0 defines whether epitope can be allergen or not.

#### 2.4.3. Toxicity determination of the T cell epitopes

ToxinPred predicts the toxicity of the epitope. In this Swiss-Prot based (SVM) prediction method was used with an E-value cut-off value of 10 for the motif-based method. SVM threshold was set as 0.0.

#### 2.4.4. Stability prediction of the T cell epitopes

ProtParam tool computes the physical and chemical parameters that are stored in the SWISS-Prot or TrEMBL like the stability of the epitope.

### 2.5. Population coverage analysis of T cell epitopes

Population coverage analysis was done to check the presence of the MHC-I molecule that binds to the T cell epitopes (as predicted by the immune database), in the human population of the SARS-CoV-2 affected countries. Because if these MHC-I molecules are present in the human population that only it will bind with the T cell epitope for eliciting an immune response in the host against SARS-CoV-2. IDEB analysis tool was used for population coverage analysis.

### 2.6. 3D structure prediction of T cell epitopes and MHC-I

As the 3D structure of the T cell epitopes and the MHC-I molecules were not present in the database, they were 3D modeled. The T cell epitopes and MHC-I 3D structures were predicted and modeled using PEP-FOLDD and SWISS-MODEL, respectively.

### 2.7. Molecular docking analysis of T cell epitopes

Molecular docking analysis of T cell epitopes were done in order to check the binding affinities between the chosen T cell epitopes and their corresponding MHC-I molecules that were derived from the immune databases. Autodock flexible receptor (ADFR) tool was used to do molecular docking for knowing the receptor-ligand interactions and binding affinities.

### 2.8. Molecular dynamics simulation of T cell epitope

The most probable T cell epitope was selected and subjected for molecular dynamic simulations in order to know the deviations and atomic fluctuations, when the T cell epitope would bind to the MHC-I molecule. GROMACS v 2018.1 was used for the molecular dynamic simulation.

### 2.9. B-cell epitope prediction

B-cell epitopes interact with the B lymphocyte for eliciting immune response^5^. IEDB tool was used for identifying B-cell epitopes and its antigenicity using methods: Kolaskar and Tongaonkar (KT) antigenicity scale prediction, Emini surface accessibility prediction, Karplus and Schulz (KS) flexibility prediction, Bepipred linear epitope prediction and Chou Fasman (CF) beta-turn prediction.

## 3. RESULTS

### 3.1. S glycoprotein sequence retrieval

10 Sequence of S glycoprotein of SARS-CoV-2 with PDB ID: 6X2A, 6X2B, 6X2C, 6X29, 6YM0, 6YLA, 6VXX, 6VSB, 6WPT and 6WPS were derived from PDB RCSB.

### 3.2. MSA

MSA predicted the conserved region from S glycoprotein as shown in Figure 2, which starts from 289 proline and ends at 461 asparagine. The conserved S glycoprotein region of the sequence is 172 amino acid long and this peptide sequence was used in further analysis.

**Figure2:**
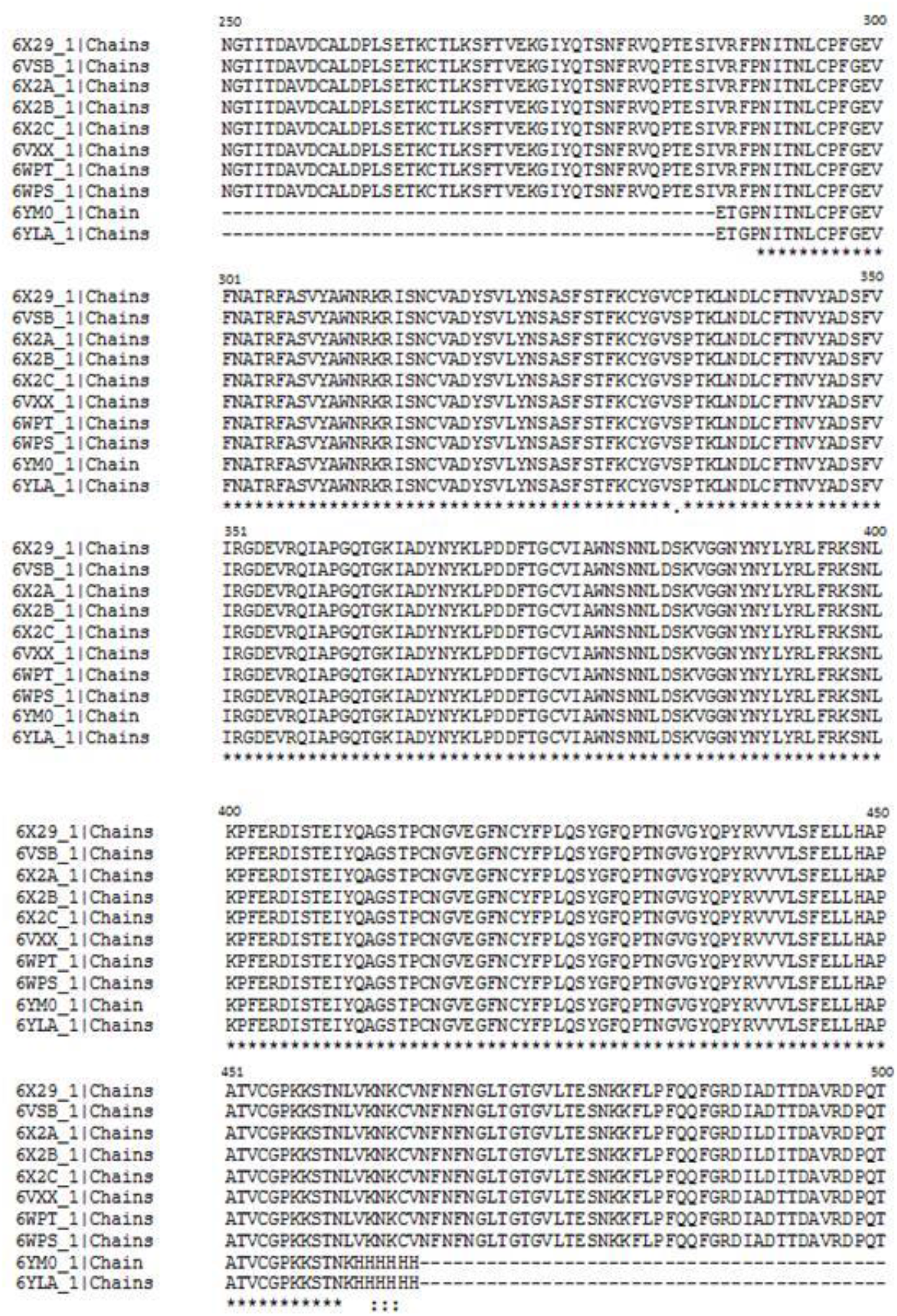
Conserved region of the spike glycoprotein (*) of SARS-CoV-2.

### 3.3. T-cell epitope prediction

From the conserved S glycoprotein region of the protein, potential T cell epitopes were taken from the NetCTL server in a preselected environment. 23 T cell epitopes were found and used for the further analysis as shown in Table 1.

**Table1:**
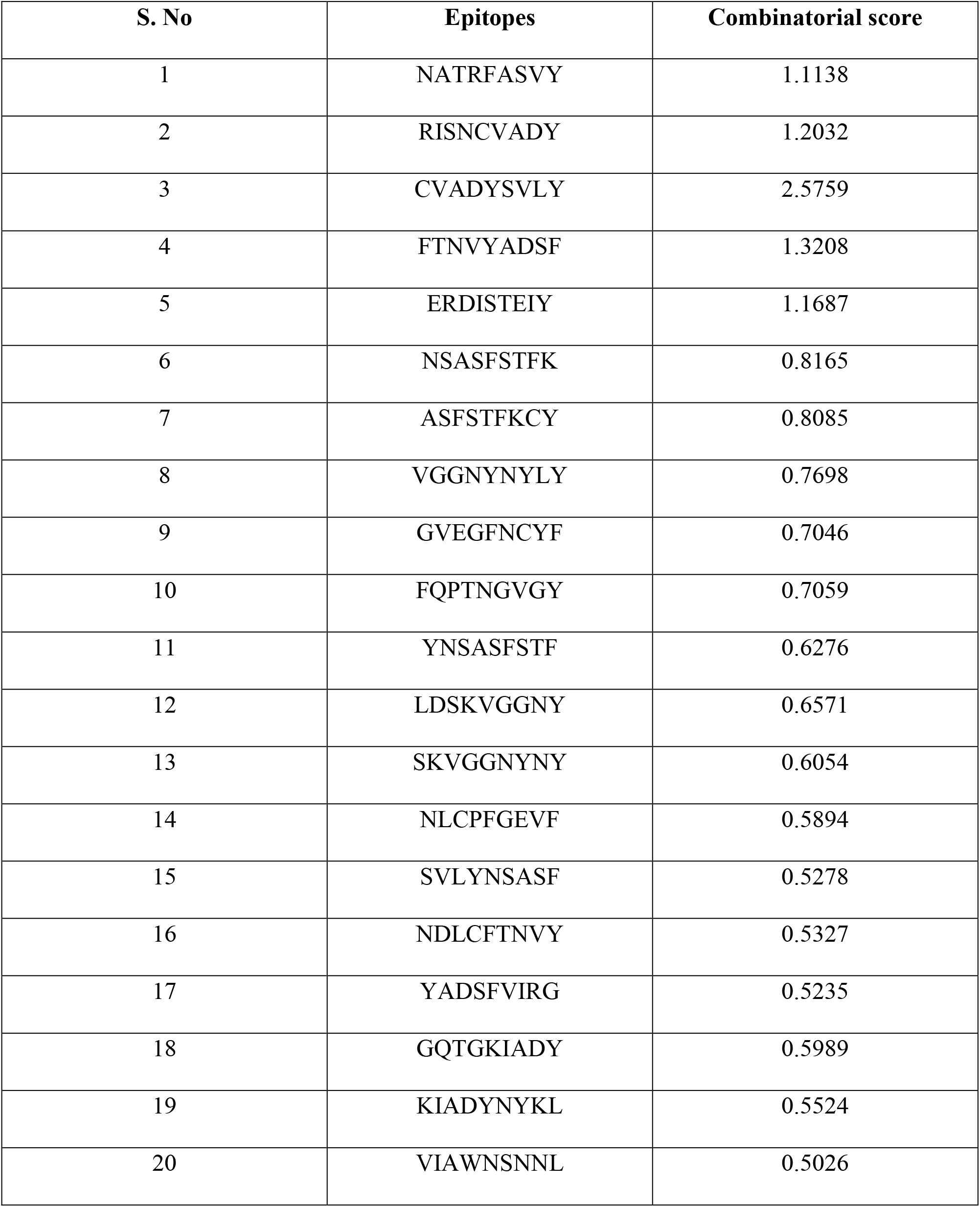
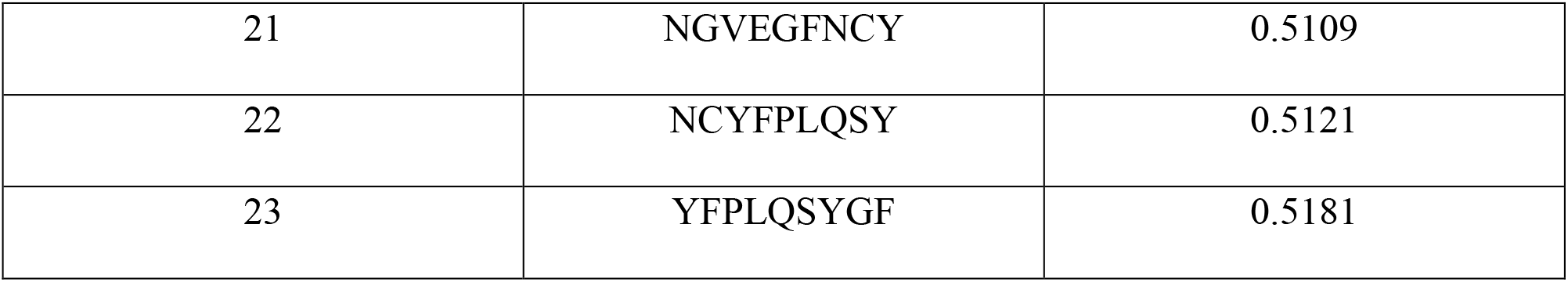
Shows the best 23 epitopes that are selected on the basis of combinatorial score.

Proteasome complex cleaves the peptide bonds and converts the proteins into small peptides. These small peptide molecules gets associates with class-I MHC molecules and are further presented to T helper cells. The binding predictions of MHC-I and processing were done from IDEB tool that generates proteasomal CT processing, TAP transport, MHC-I and processing score. The overall score predicts the peptides potential to be a T-cell epitope as shown in Table 2.

**Table2:**
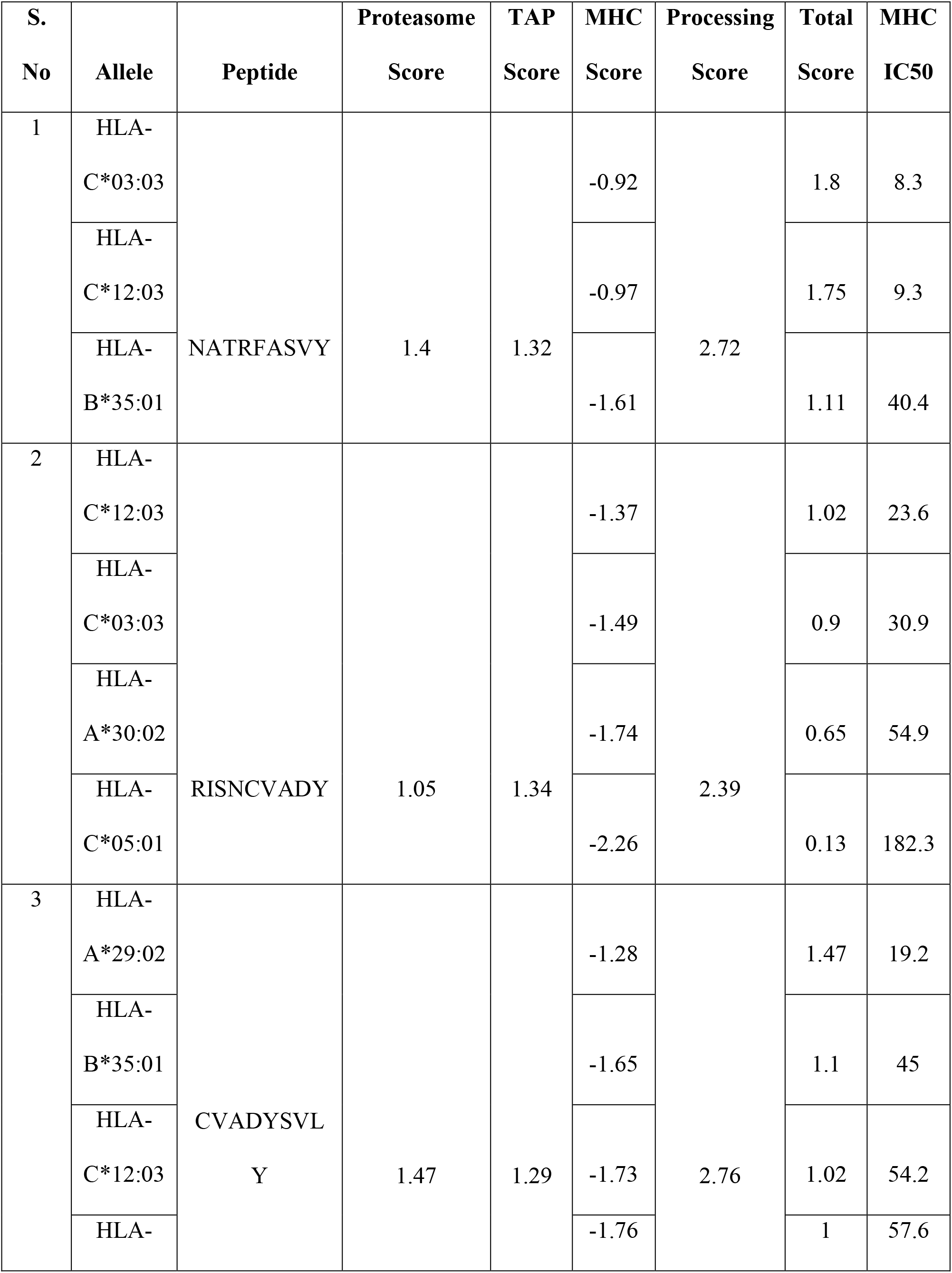
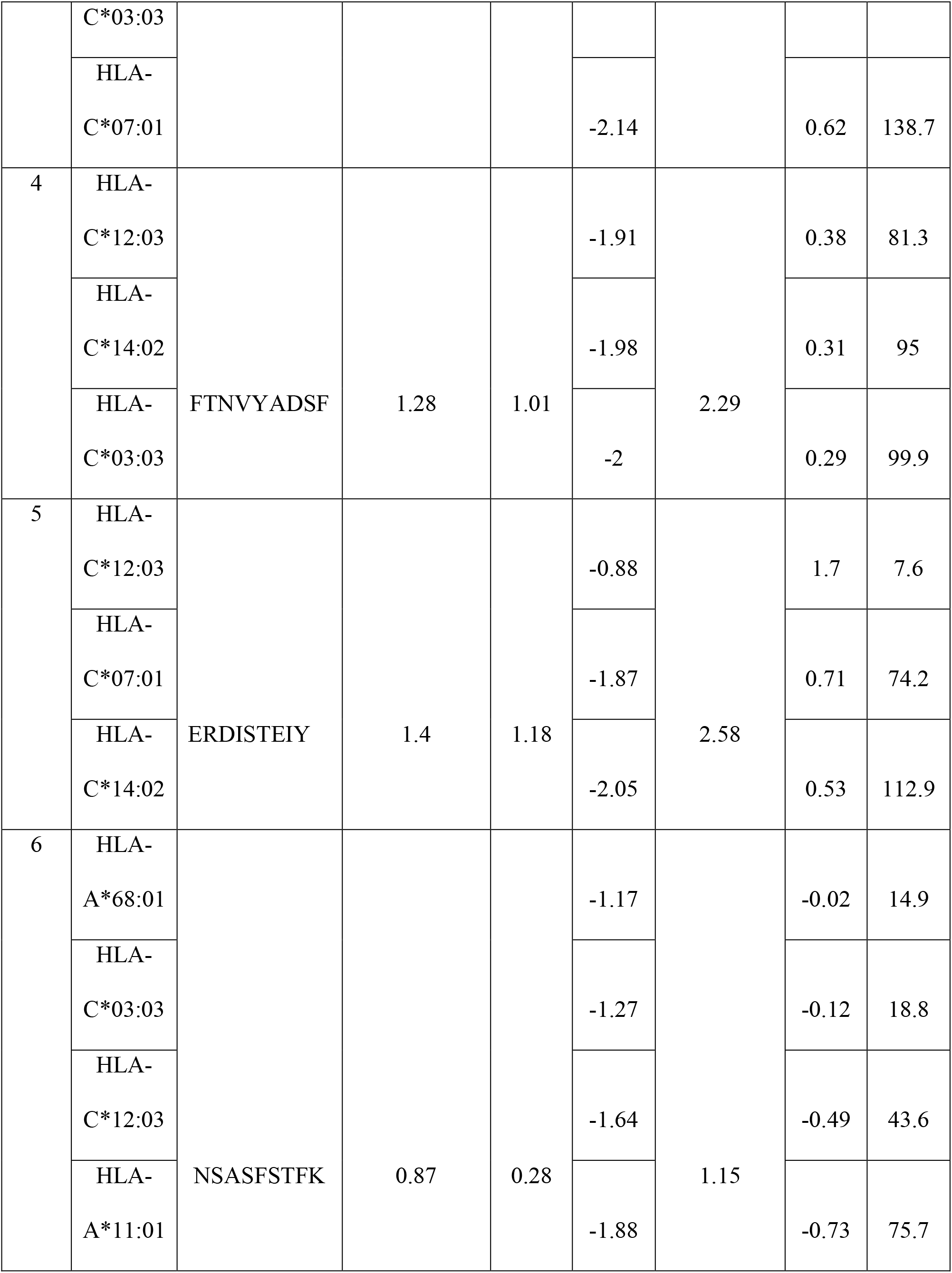
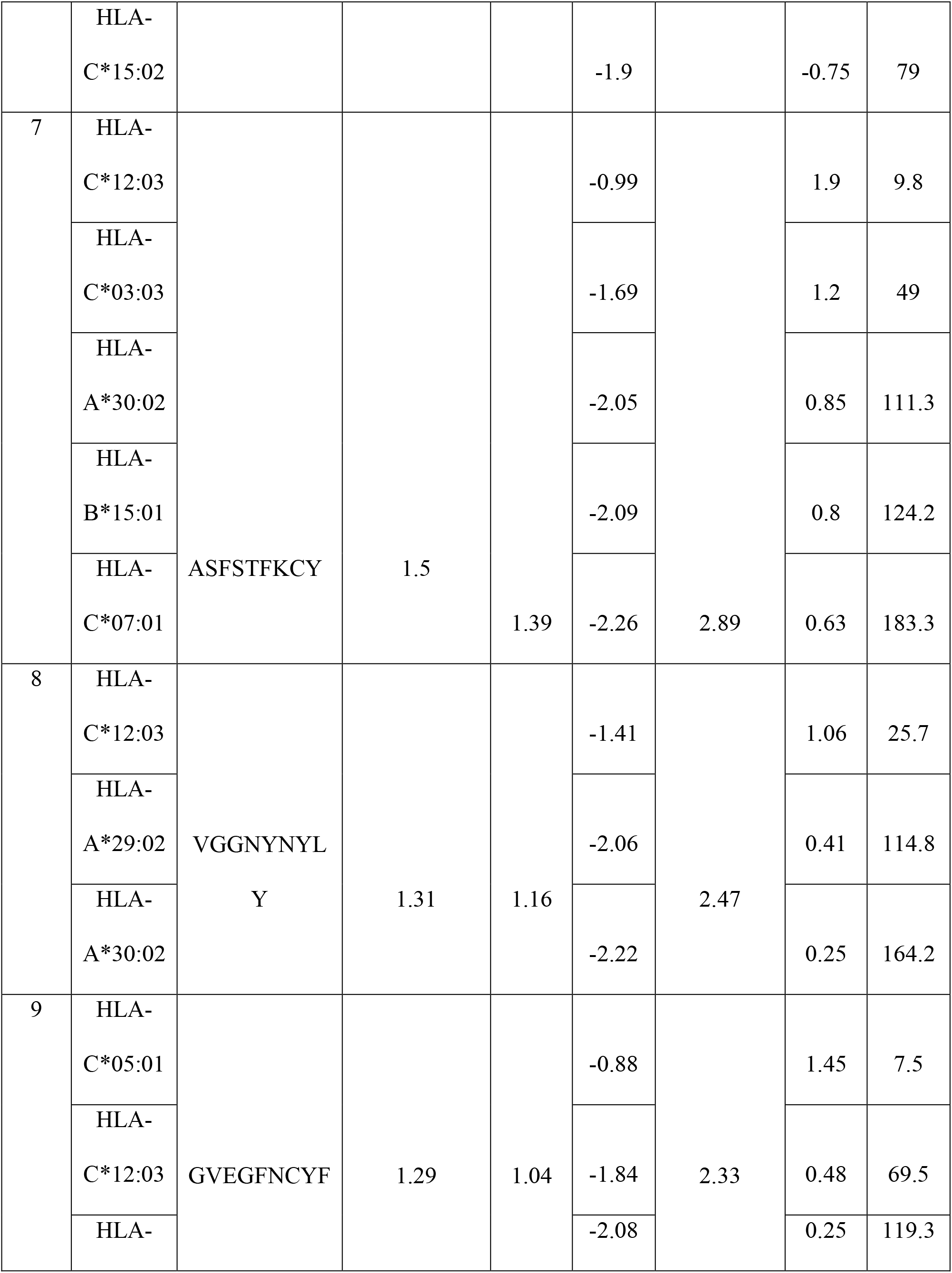
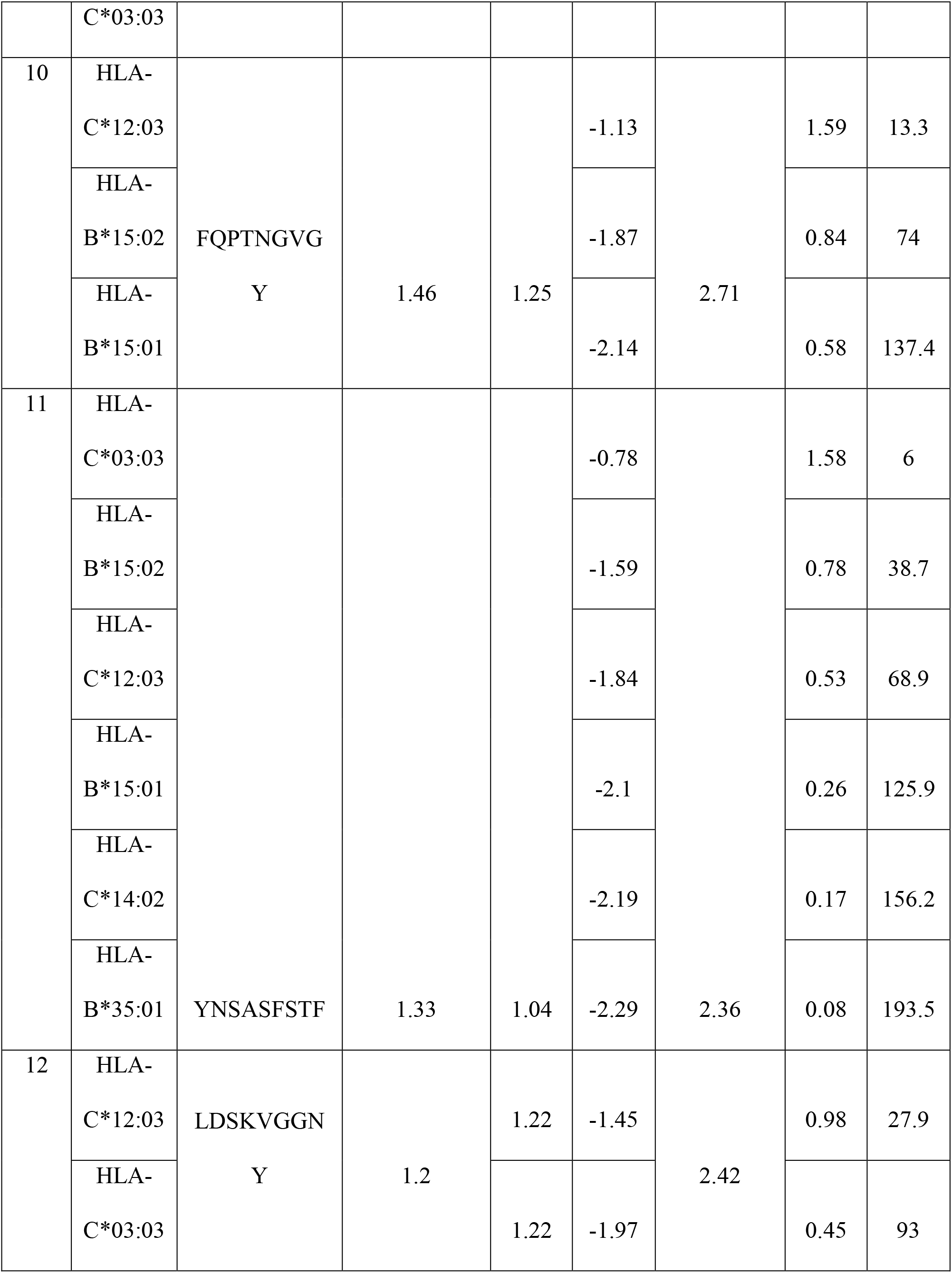
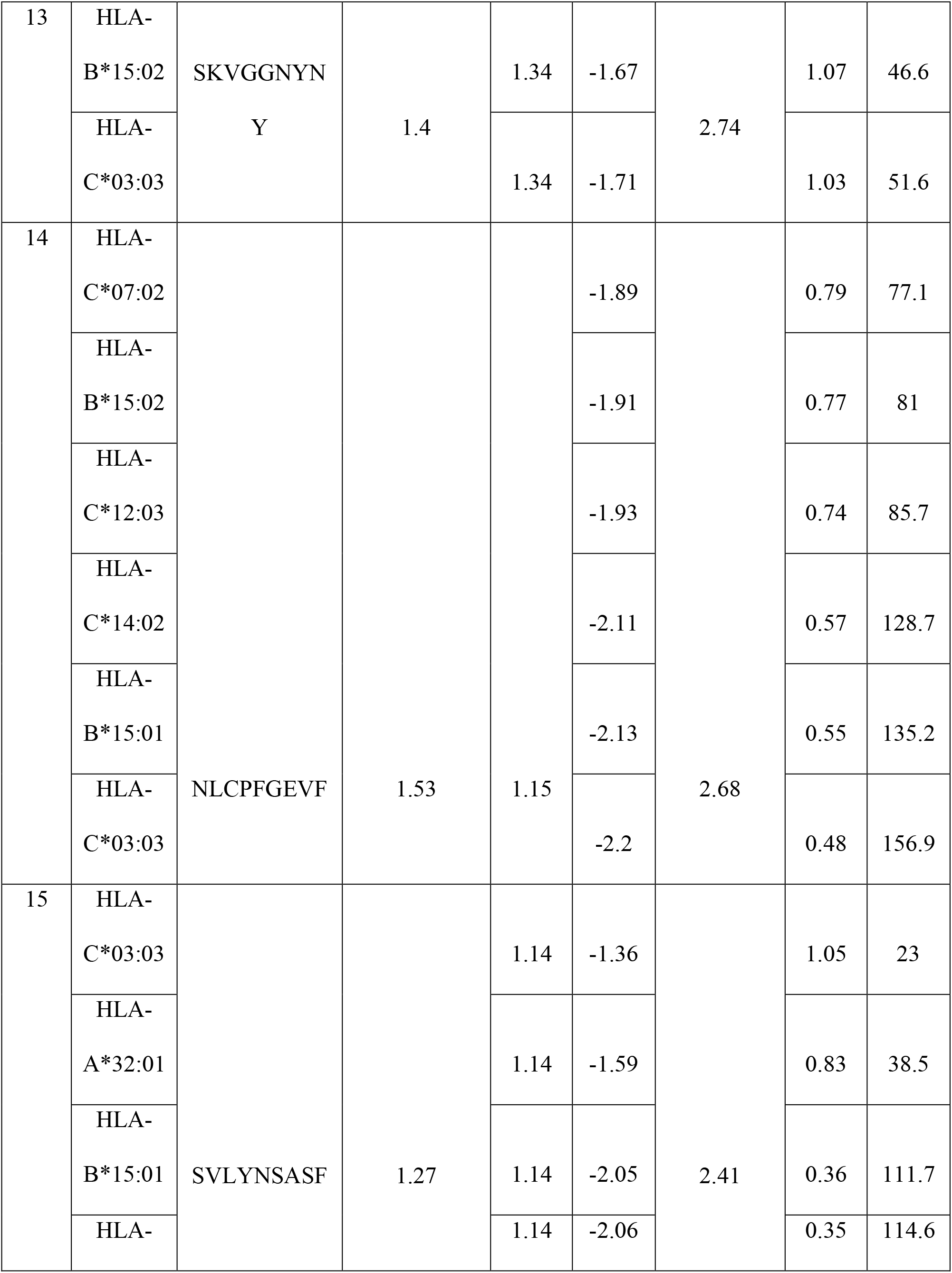
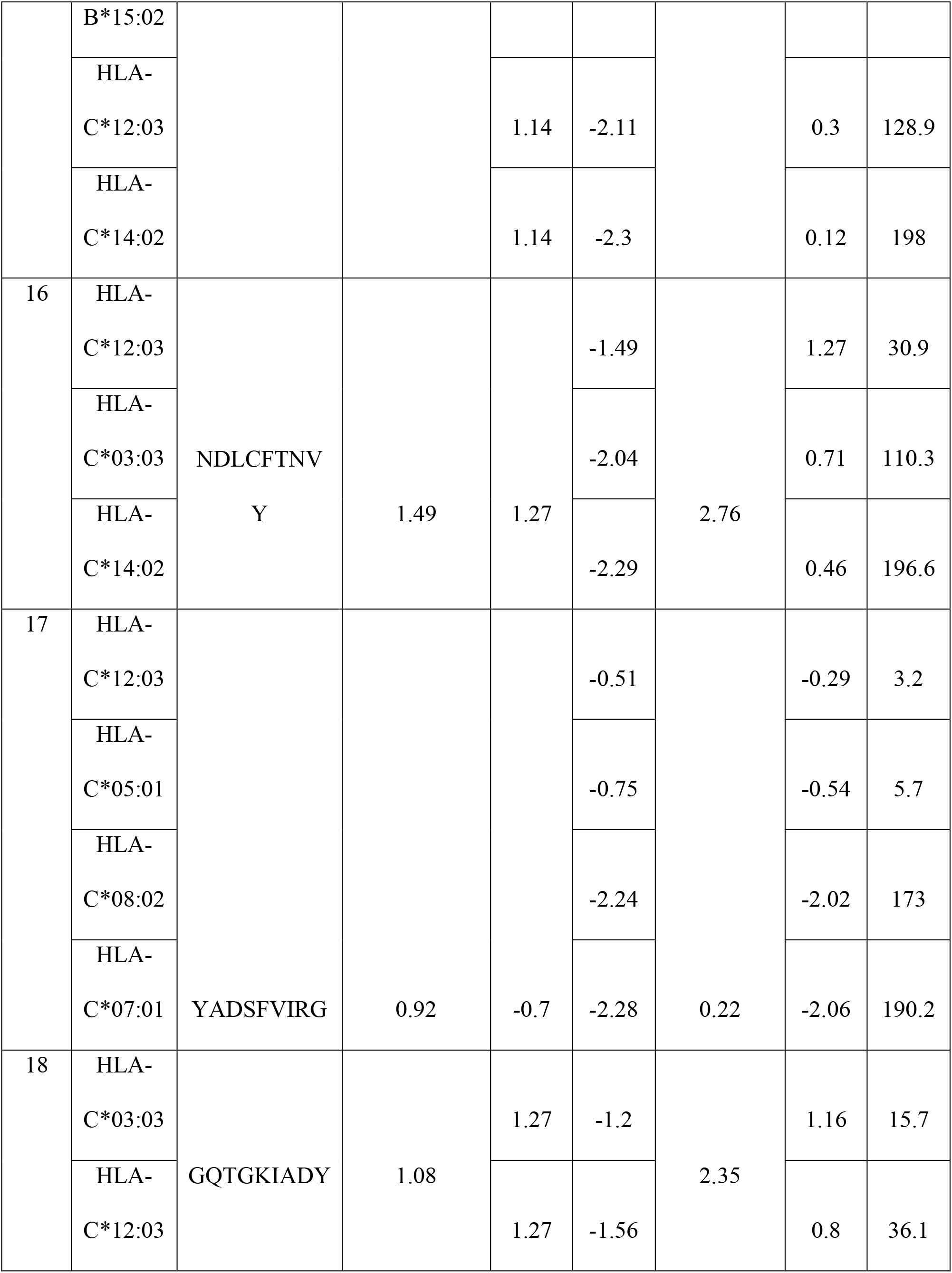
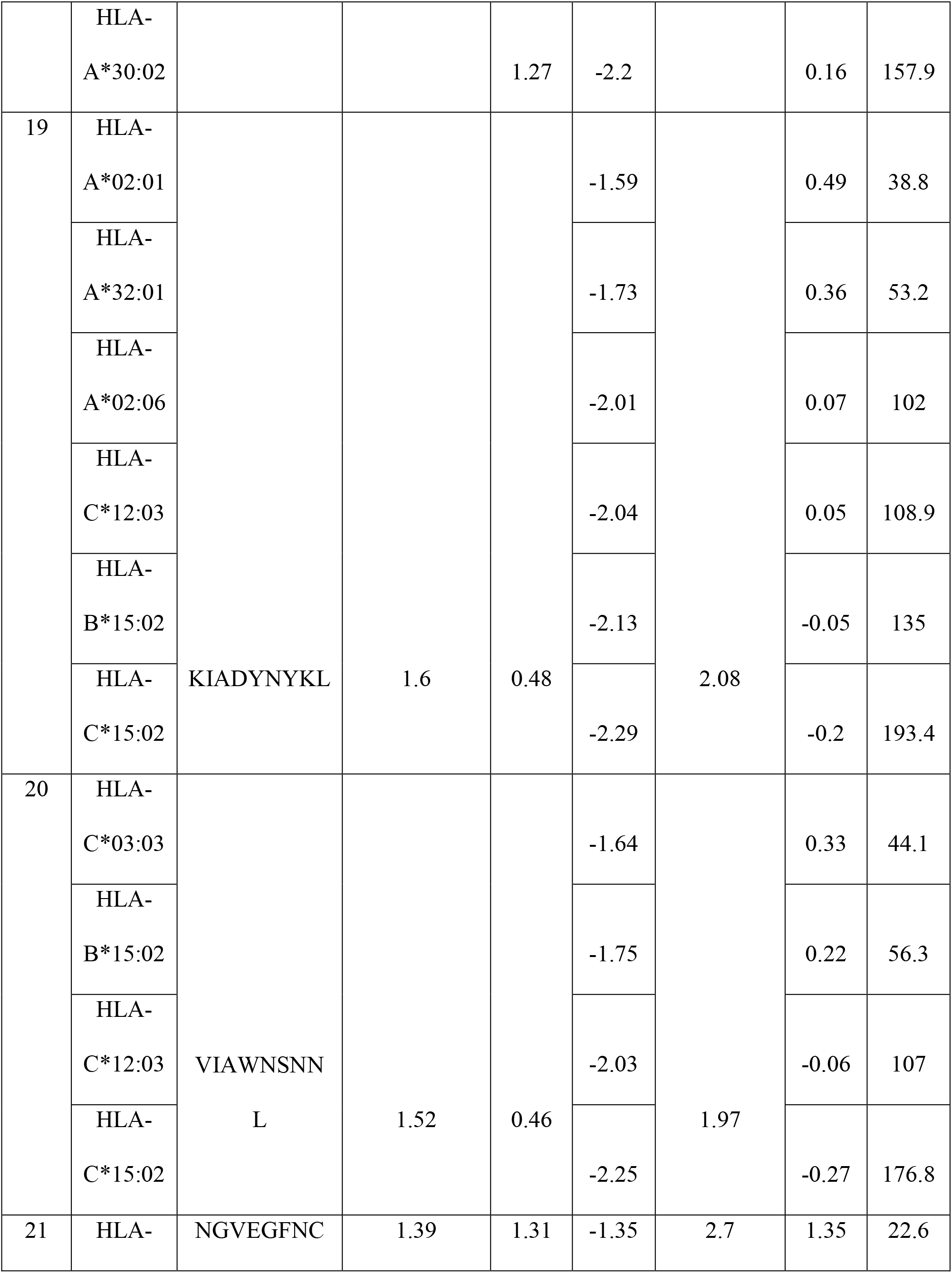
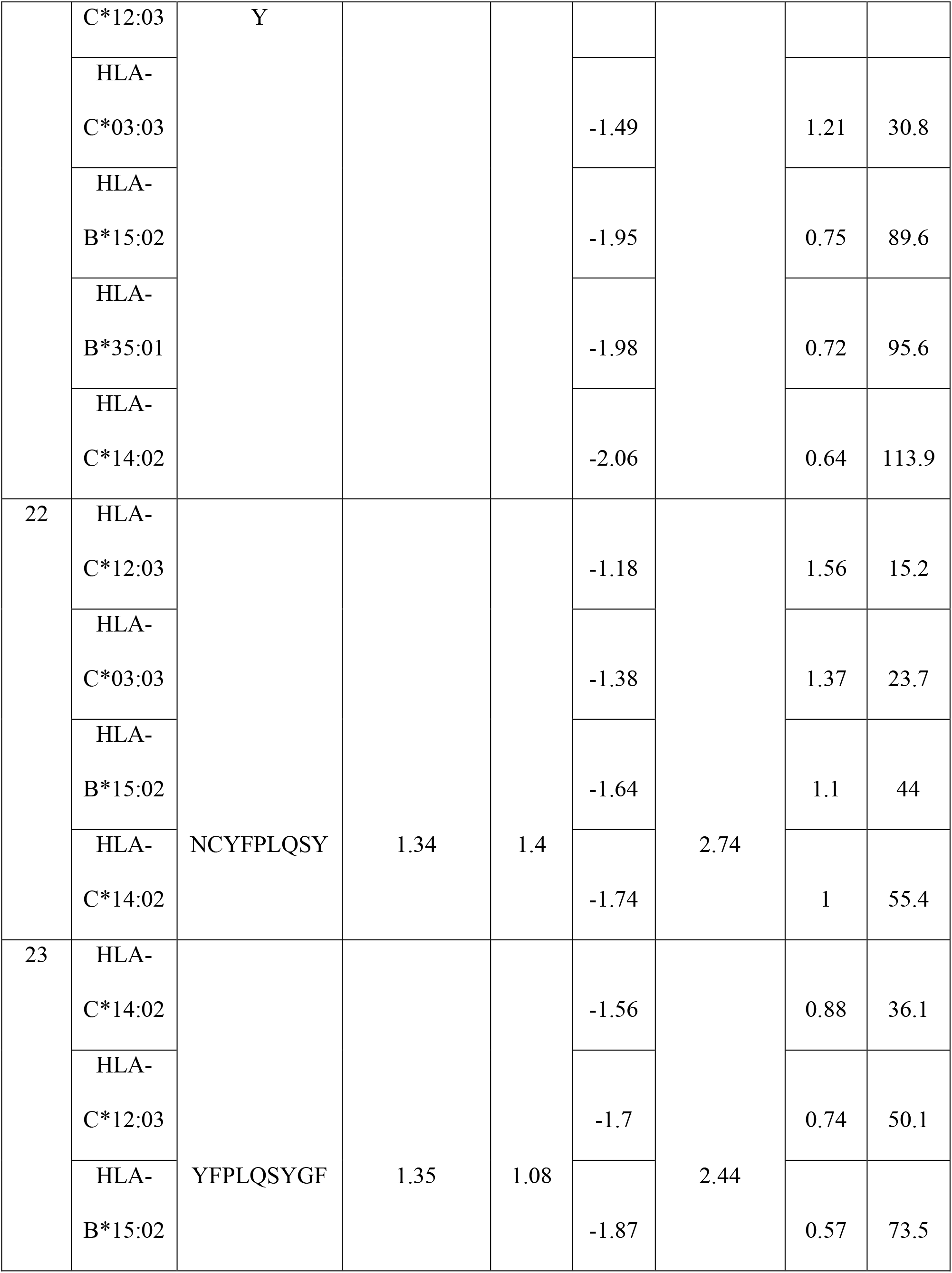
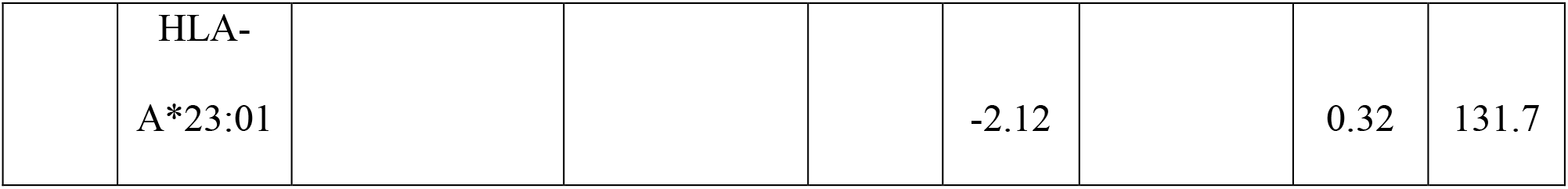
23 potential T cell epitopes along with their interacting MHC-I alleles, proteasomal cleavage score, TAP transport score, MHC score, processing score and the overall total score that predicts the peptides intrinsic potential to be a potential T cell epitope that could elicit immunogenic response in the host.

MHC-I binding predictions resulted in a range of MHC-I alleles that interacts with selected T cell epitopes. The MHC-I that were having highest binding affinity were chosen for further analysis.

### 3.4. Physiochemical analysis of T cell epitopes

For an epitope to be effective and safe for the host, antegenicity, non-allergenicity, non-toxicity and stability are given some parameters on the basis of which 4 epitopes were selected for further analysis (LDSKVGGNY, SKVGGNYNY, NDLCFTNVY and GQTGKIADY) as shown in Table 3.

**Table 3:**
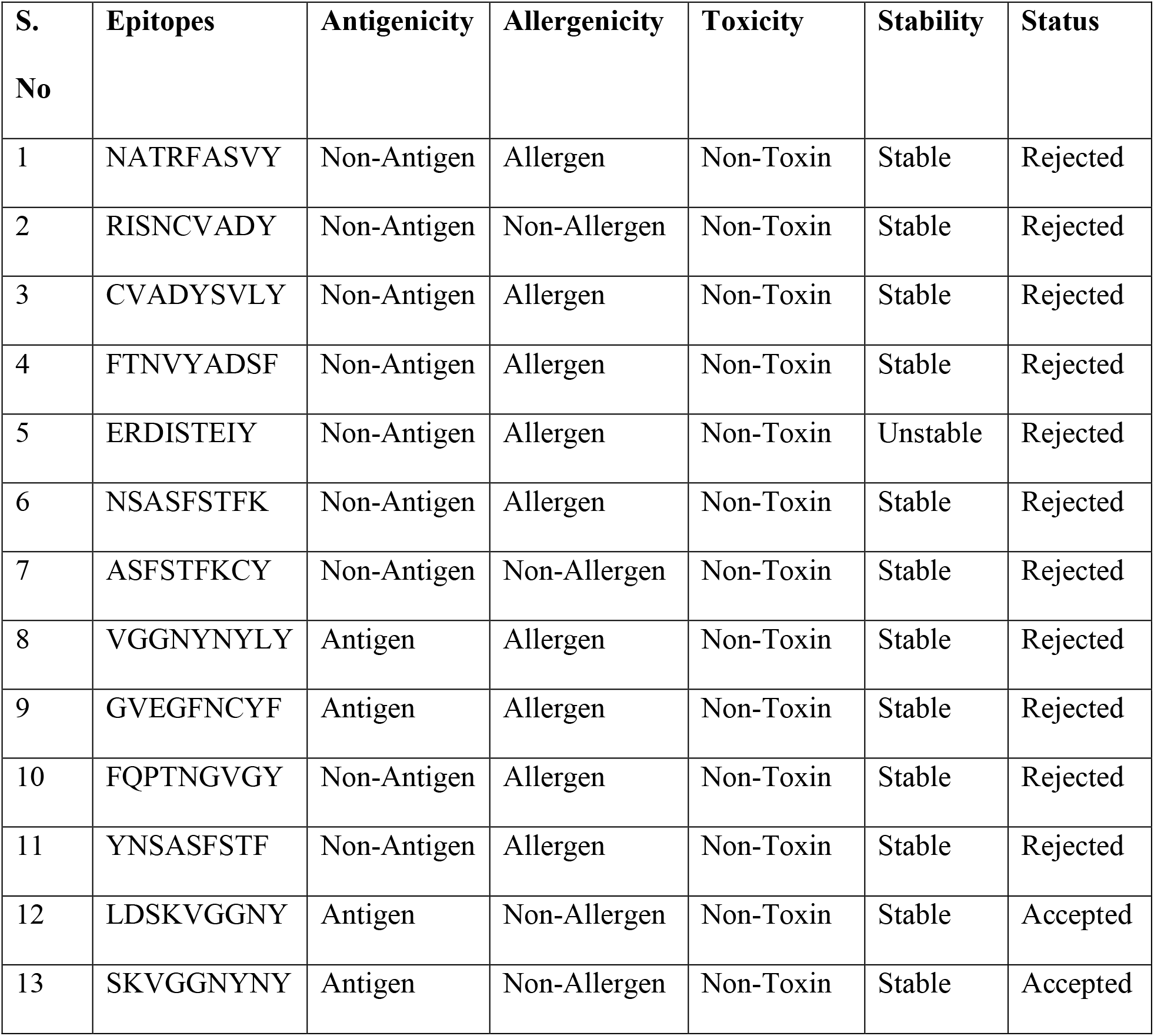
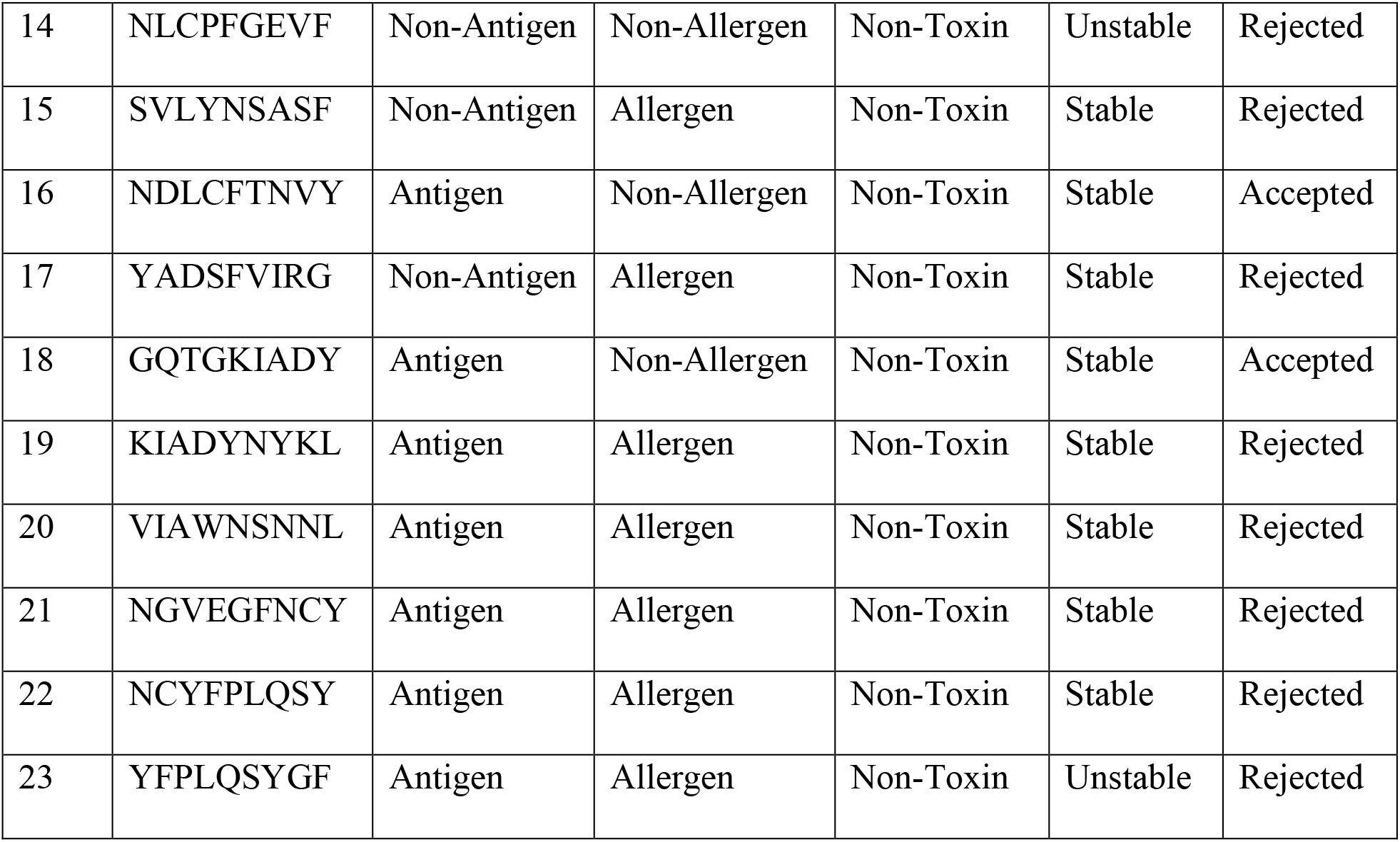
Shows the physiochemical properties of the epitope.

### 3.5. Population coverage analysis of T cell epitopes

The population coverage of the predicted epitopes is depicted in Table 4. T-cell recognizes a complex of MHC and pathogen-derived epitope. This means that the particular MHC needs to be present in the individual, so that it can binds to a particular T-cell pathogen-derived epitope for eliciting an immune response. This is known as MHC restriction of T-cell response. As MHC molecules are polymorphic and different human leukocyte antigen (HLA) alleles are present in human population. If peptides are selected that binds with HLA with a high affinity and that HLA is present in target human population, then epitope-based vaccine could be more effective. Therefore, careful considerations must be taken care so that the vaccine is not ethnically biased. For the issue discussed above, IEDB population coverage analysis helps in calculating the fraction of individuals that contains the predicted MHC.

**Table 4:**
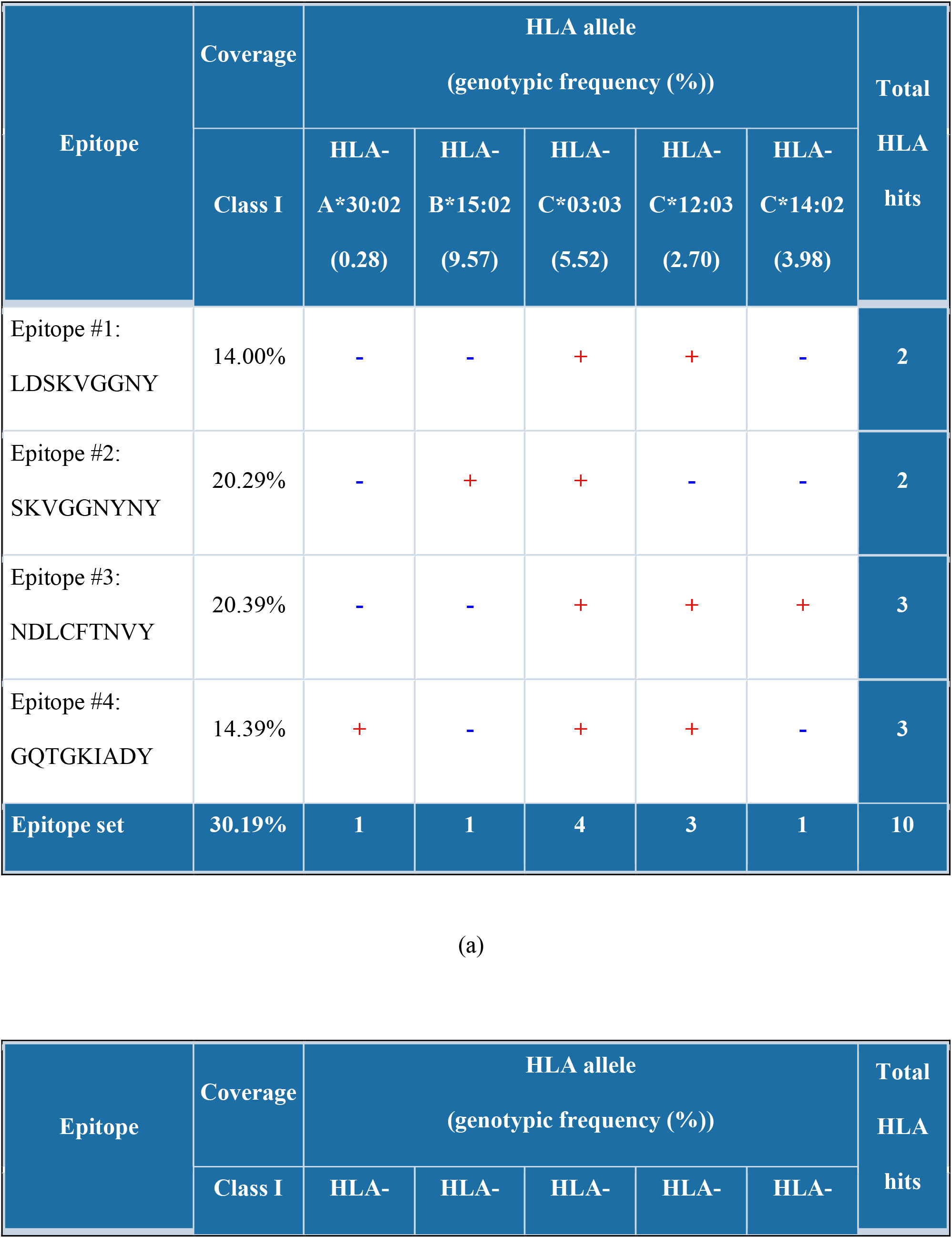
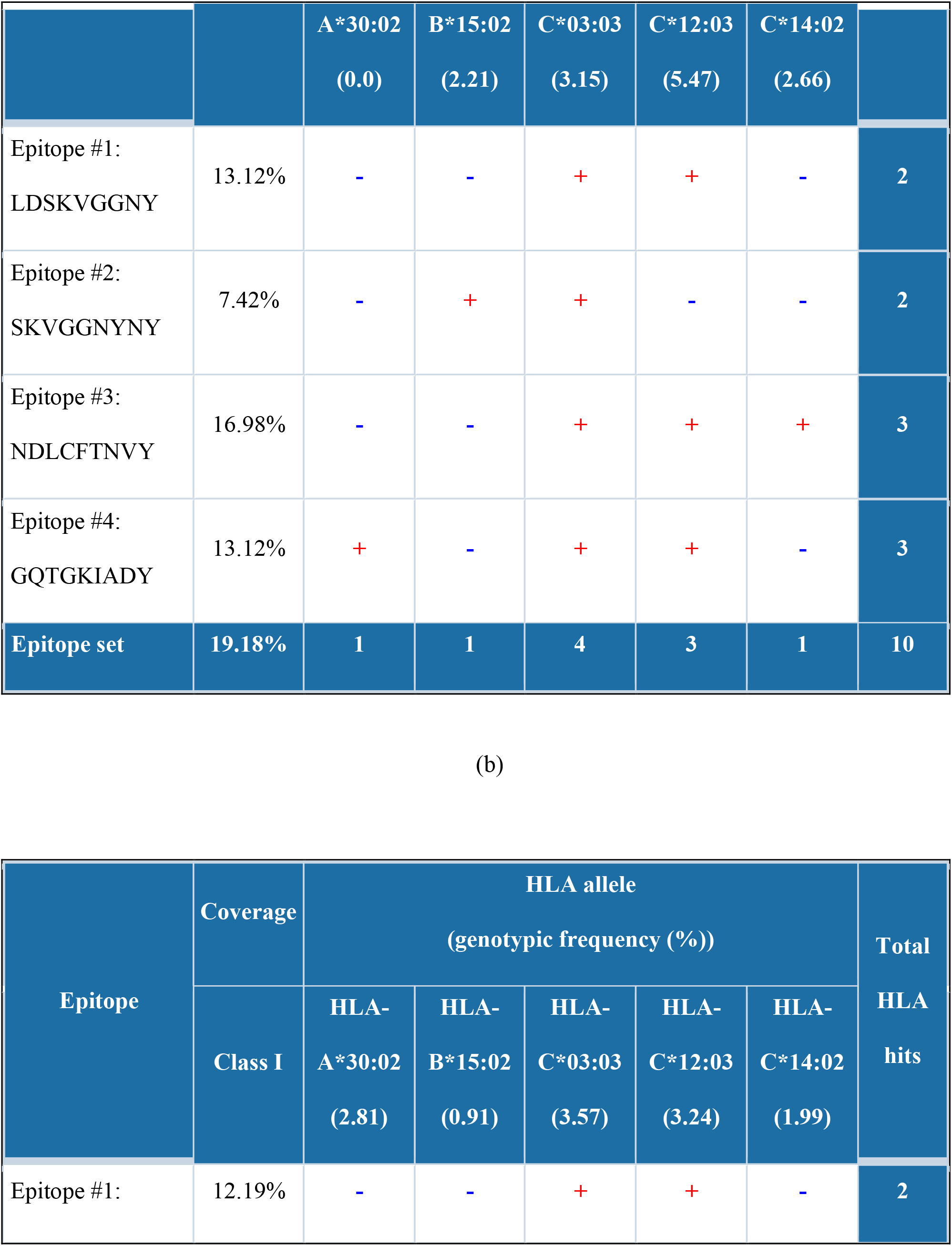
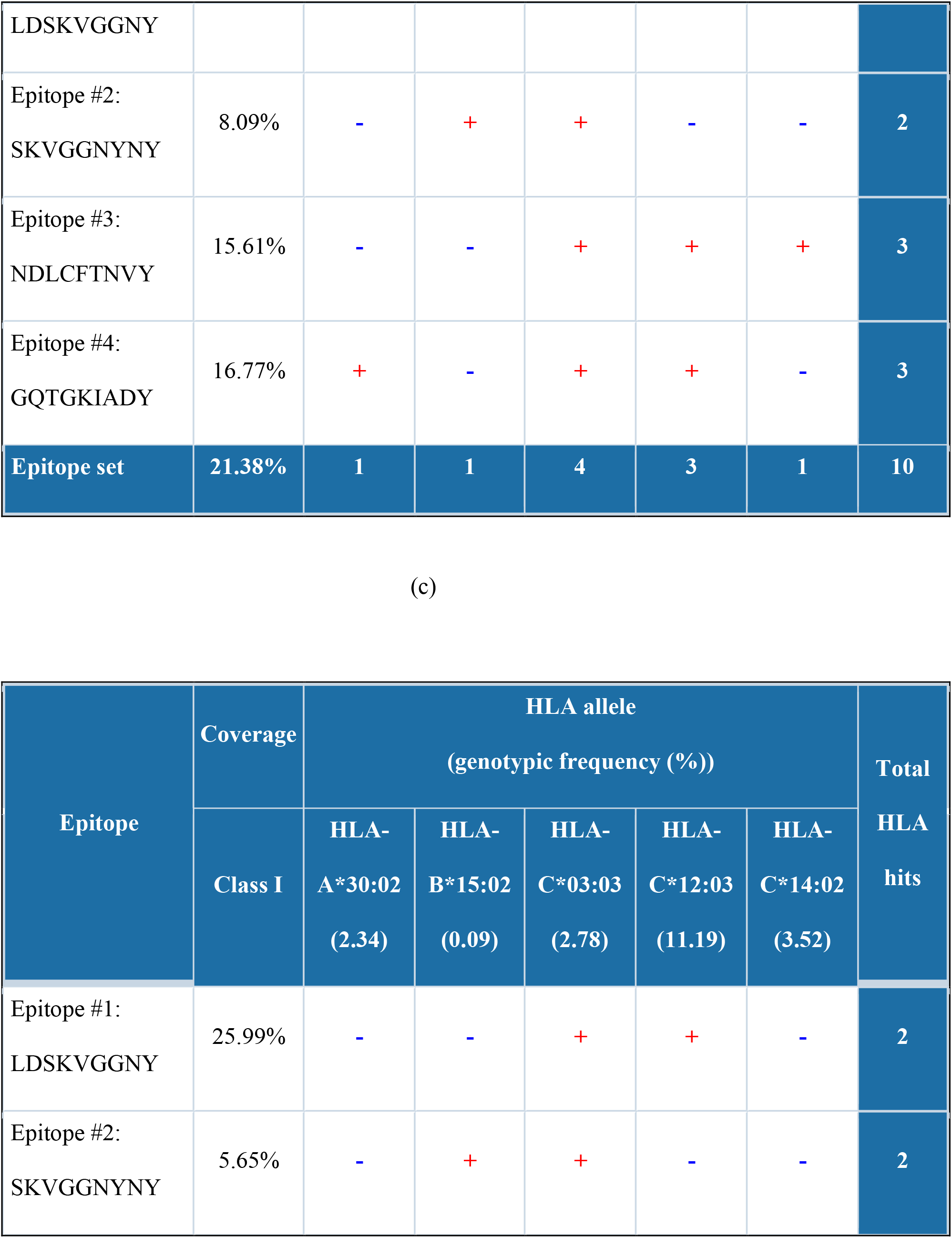
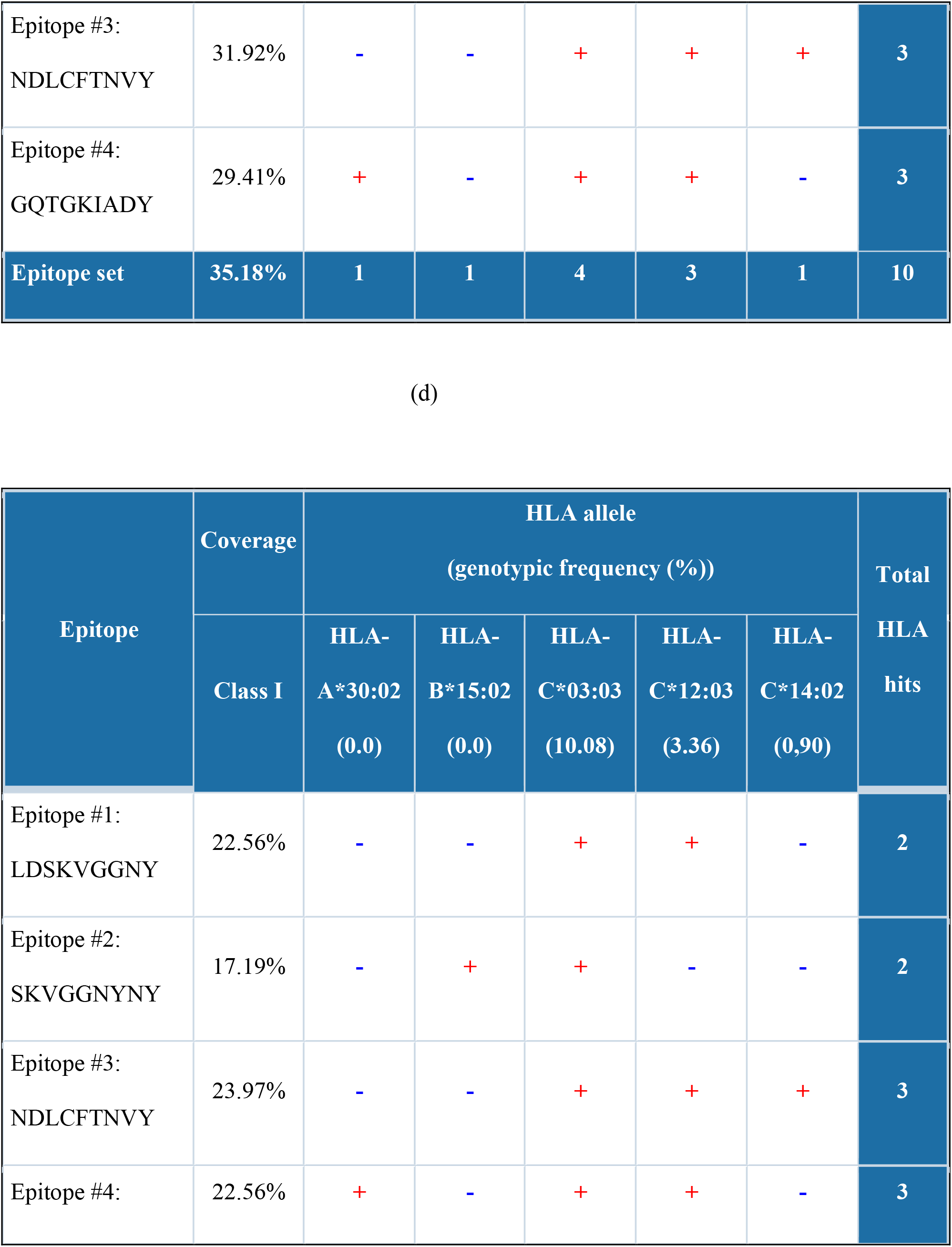
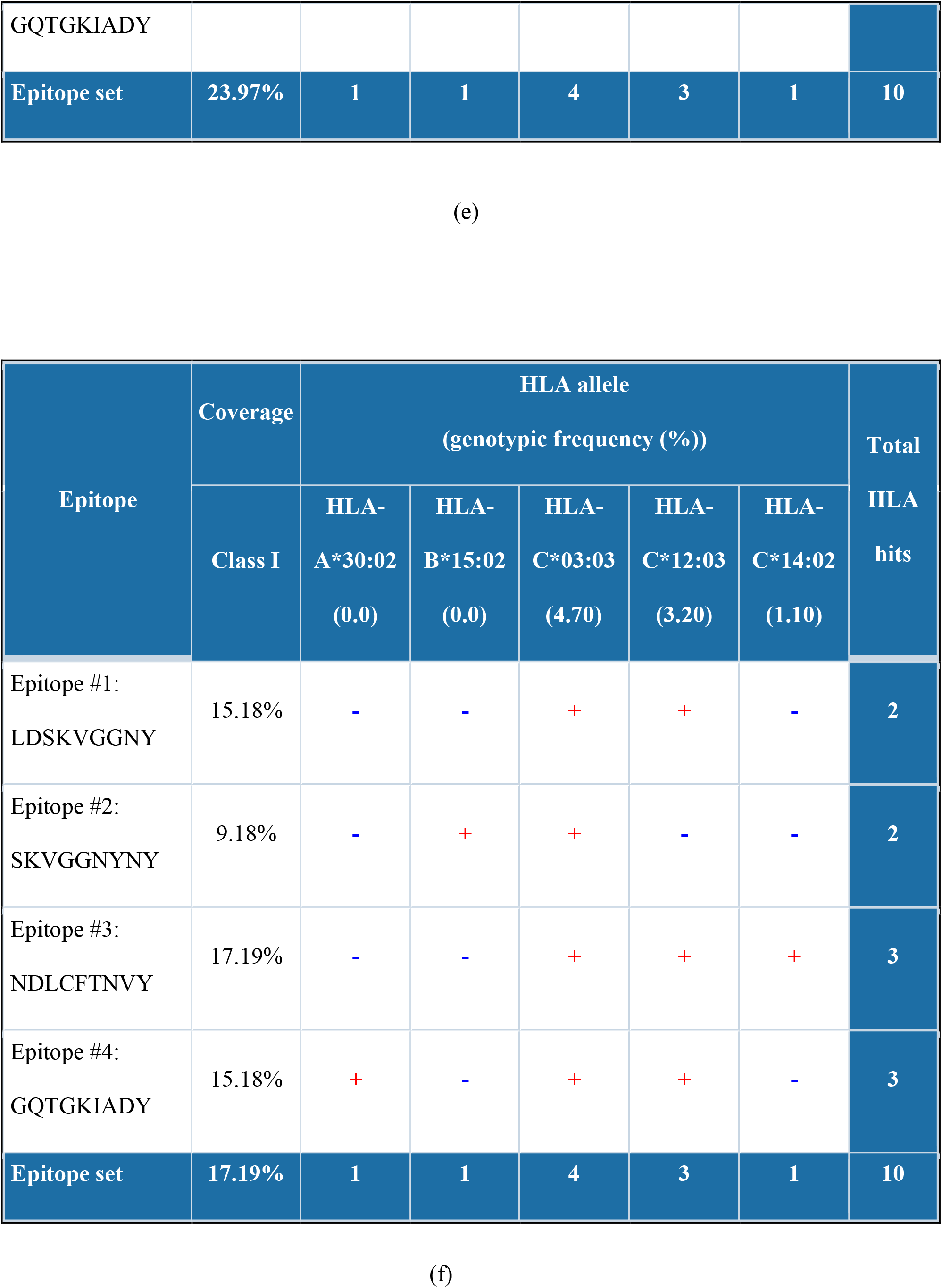
Population coverage of chosen epitope shown based on MHC-I restriction data in most effected countries (a) China (b) India (c) United States (d) Italy (e) Russia (f) the United Kingdom. (+) means MHC-I restricted and (−) means not MHC-I restricted.

### 3.6. 3D structure prediction of T cell epitopes

3D structure modeling of T cell epitopes were done using homology modeling approach using SWISS MODEL web server that is fully automated and freely available. Further, MHC-I were modeled using a de-novo approach that predicts 3D structure of protein from its linear amino acid sequence.

### 3.7. Molecular docking analysis of T cell epitopes

ADFR tool was used to find the binding affinities between the T-cell epitope and MHC-I that is having a lower value of IC_50_. GQTGKIADY T-cell epitope binds with HLA-C*03:03 most strongly as described in Table 5 and Figure 3, as it is having the highest binding affinity.

**Table 5:**
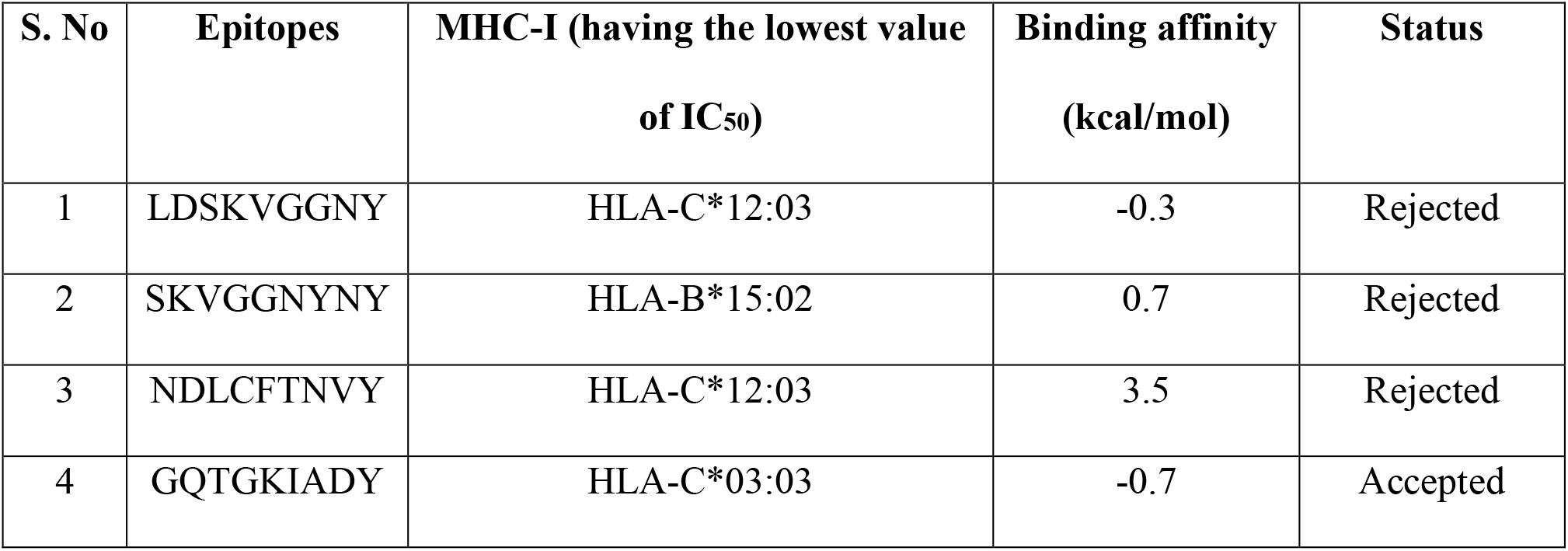
Shows the binding affinities between the 4 selected T cell epitopes and the MHC-I that have lowest value of IC_50_

**Figure 3:**
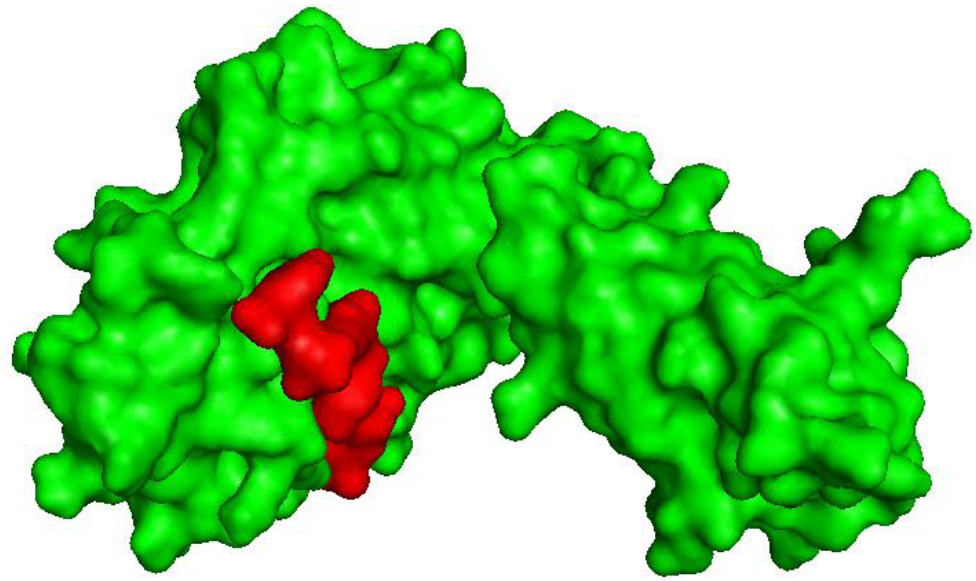
Shows surface interaction of GQTGKIADT T cell epitope (red) with HLA-C*03:03 (green).

### 3.8. Molecular dynamic (MD) simulation of T cell epitope

MD simulations were applied for the modeled structure of T-cell epitope GQTGKIADT and HLA-C*03:03 complex. This was done in order to understand the stability and dynamics of the complex for 50 ns in GROMACS. The system was solvated in TIP3P salvation box and CHARMM36 all atom force field.

Root mean square deviation (RMSD) was performed to calculate the average distance of the backbone C-alpha (C_α_) atom of the superimposed frames as observed in Figure 4a. An initial change can be observed between 0 to 10 ns. After that, another change is observed between 30 to 40 ns. After which the system gets quite stable. Next, Root mean square fluctuation (RMSF) was applied to the system trajectories as observed in Figure 4b. RMSF calculates the average residual mobility of complex residues from its mean position. Minor fluctuations can be observed, which mean that the complex is stable, except between 30 to 40 ns. As observed from the Figures 4a and 4b, the complex of T-cell epitope GQTGKIADT and HLA-C*03:03 is stable after 40 ns.

**Figure 4:**
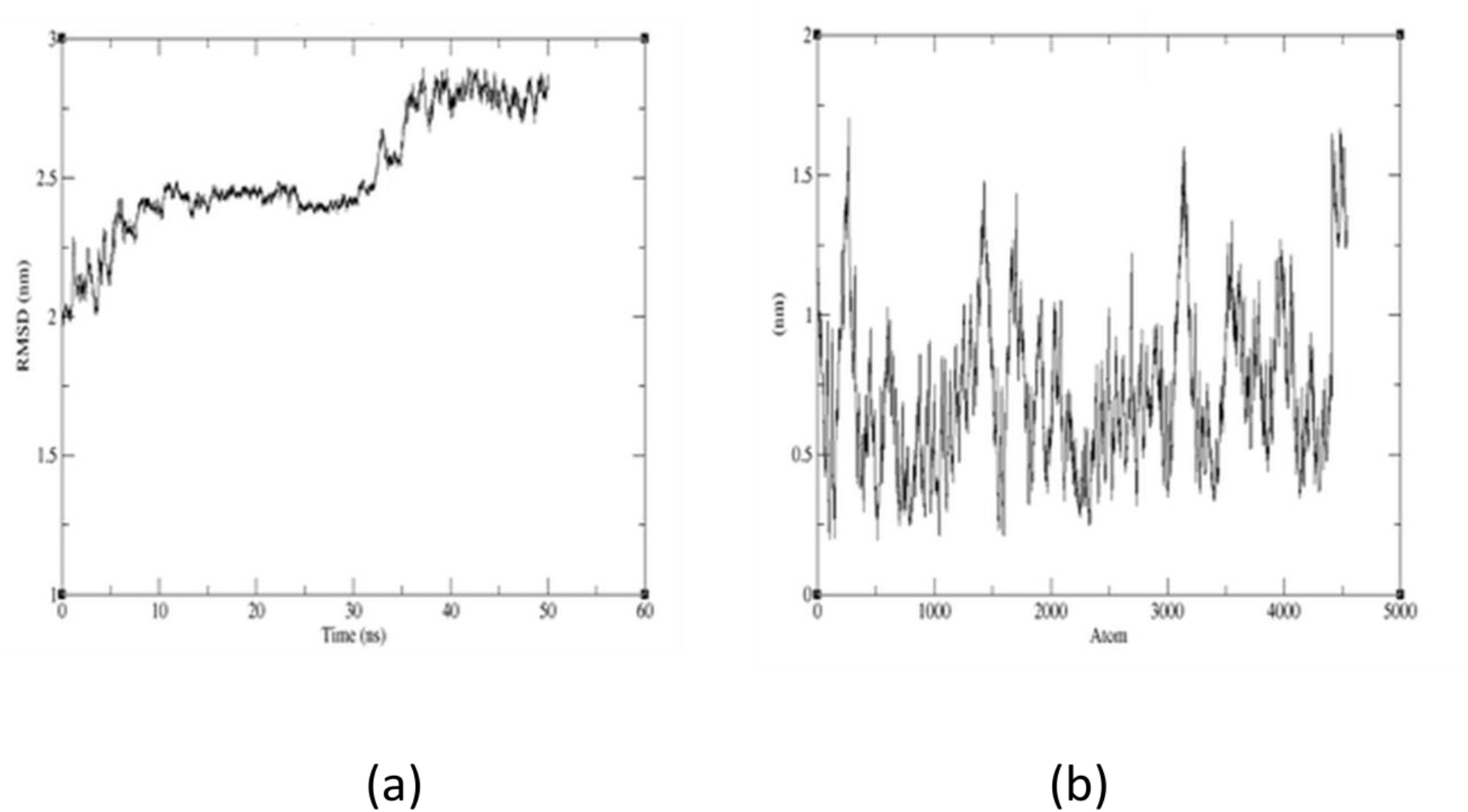
(a) RMSD and (b) RMSF plot of T cell epitope GQTGKIADT and HLA-C*03:03 complexes.

### 3.9. B-cell epitope identification

The different analysis method was used from IEDB tool to identify B-cell epitope. This tool uses amino acid scale based method.

#### 3.9.1. Prediction method of KT antigenicity

The determination of antigenicity was on the basis of physicochemical properties of amino acid. The average antigenic propensity of the conserved S glycoprotein of SARS-CoV-2 was 1.043, with a maximum value of 1.214 and minimum value of 0.907. The antigenic determination threshold was 1.043 (>1.00 are potential antigenic determinants). 6 epitopes satisfy the threshold value and so they have the capacity to express B-cell response. Results are summarized and shown in Figure 5 and Table 6.

**Figure 5:**
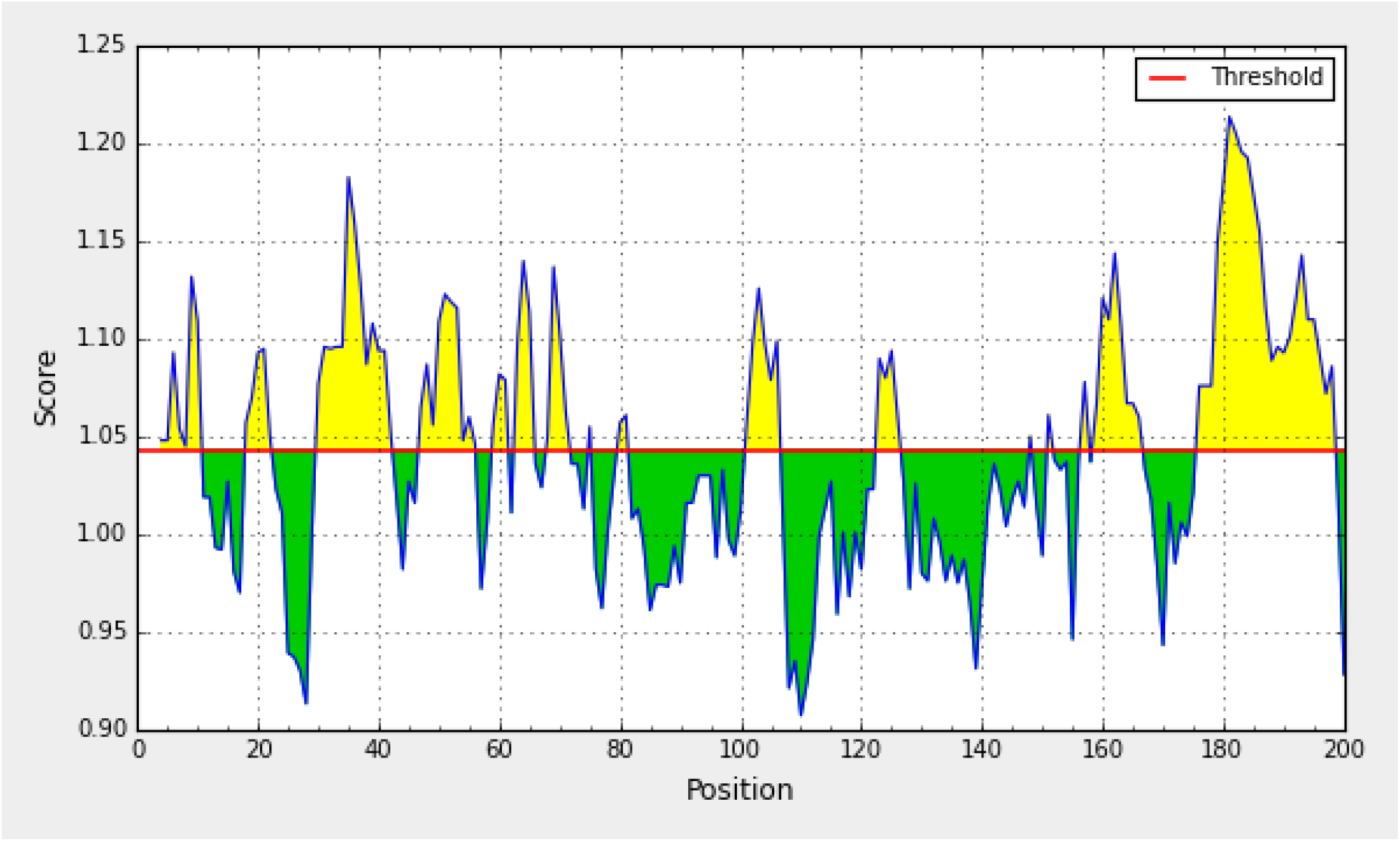
Kolaskar and Tongaonkar antigenicity analysis of the conserved region of spike glycoprotein of SARS-CoV-2.

**Table 6:**
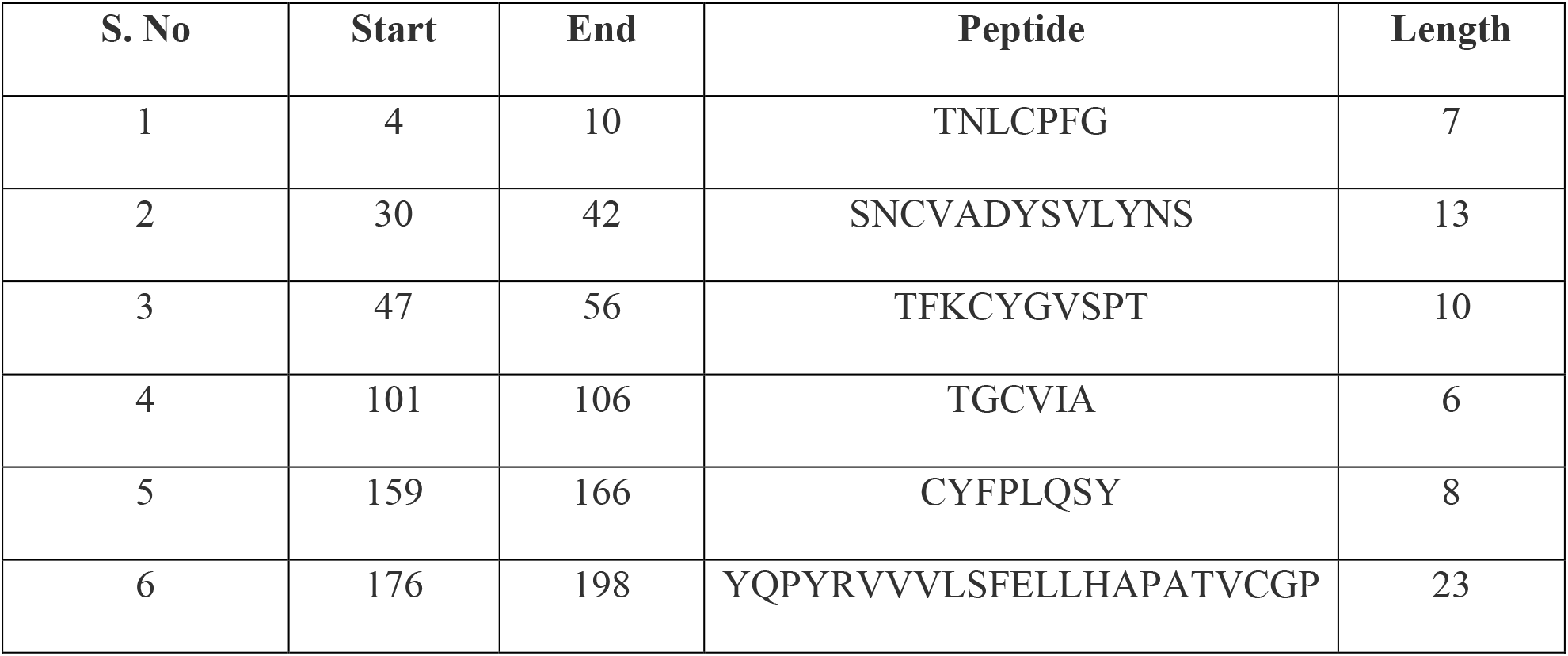
Shows B cell epitopes that have the potential to elicit an immune response, as predicted from Kolaskar and Tongaonkar antigenicity analysis of the conserved region of spike glycoprotein of SARS-CoV-2.

#### 3.9.2. Prediction method of Emini surface accessibility

For being a potential B-cell epitope, it must have surface accessibility. Therefore, this method is used to predict the peptide surface accessibility. The average value of peptide antigenic propensity was 1.00, with a maximum and minimum value of 4.805, 0.073, respectively. The antigenic determination threshold was 1.00. The region between 90 to 100 amino acid residues was found to be more accessible in the conserved S glycoprotein of SARS-CoV-2 as shown in Figure 6 and Table 7.

**Figure 6:**
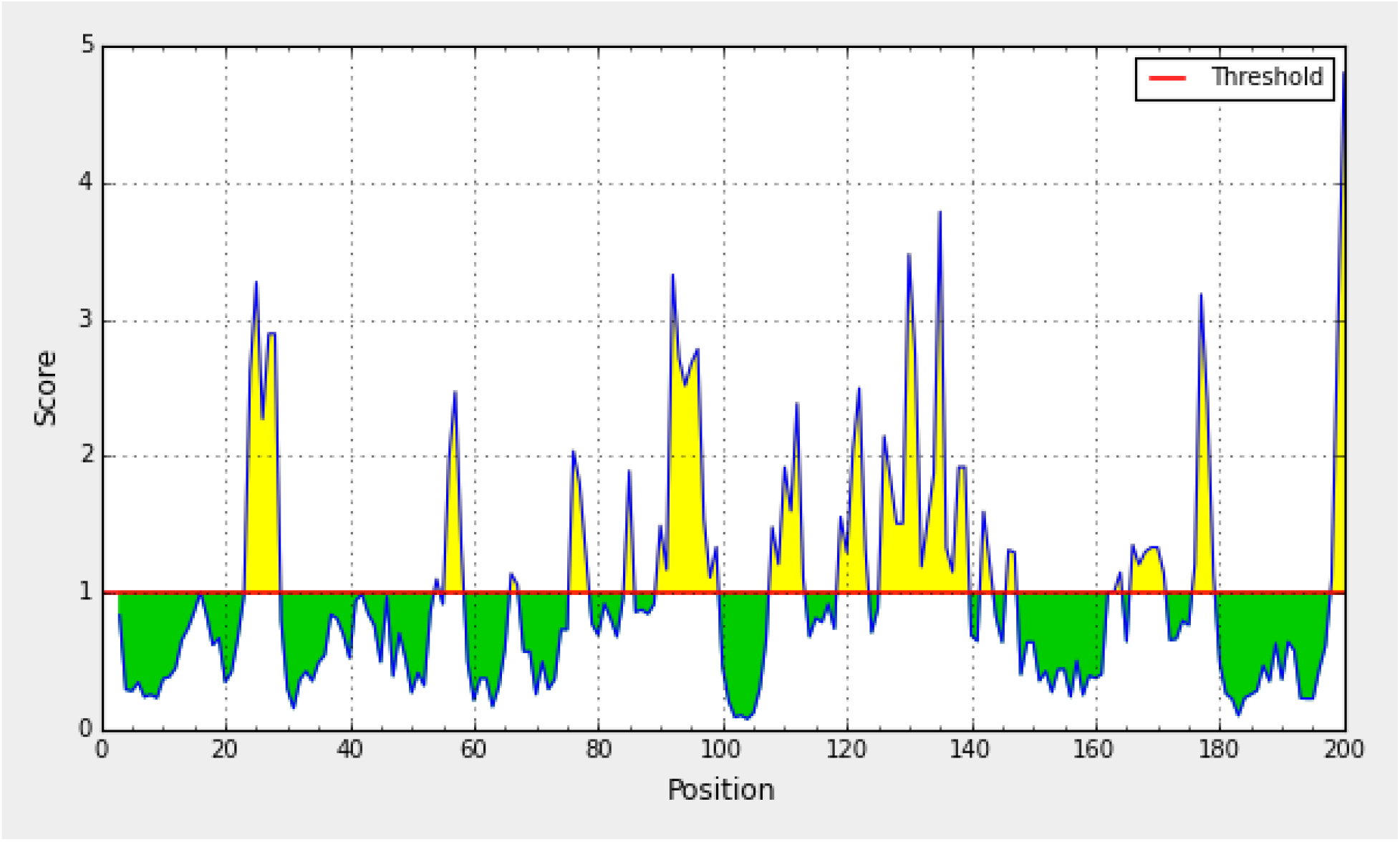
Emini surface accessibility prediction of a conserved region of spike glycoprotein of SARS-CoV-2.

**Table 7:**
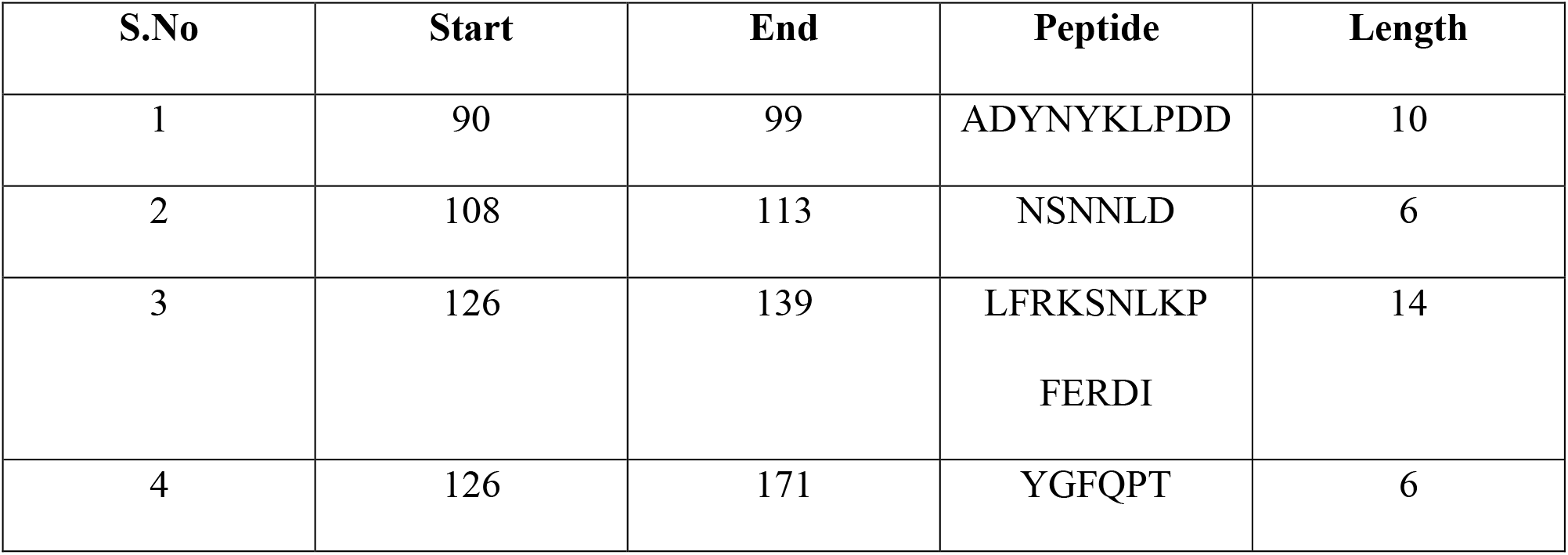
Shows B cell epitopes that are accessible at the surface, as predicted from Emini surface accessibility analysis of the conserved region of spike glycoprotein of SARS-CoV-2.

#### 3.9.3. Prediction metion of KS flexibility

Experimentally, the antigenicity is correlated with its peptide flexibility^6^. Therefore, this method was implemented to investigate the flexibility of the peptide. The average value of peptide antigenic propensity has been found 0.989, with a maximum and minimum value of 1.112 and 0.896, respectively. The threshold value of antigenic determination was found to be 0.989. The region from 80 to 88 amino-acid was found to be the most flexible as shown in Figure 7.

**Figure 7:**
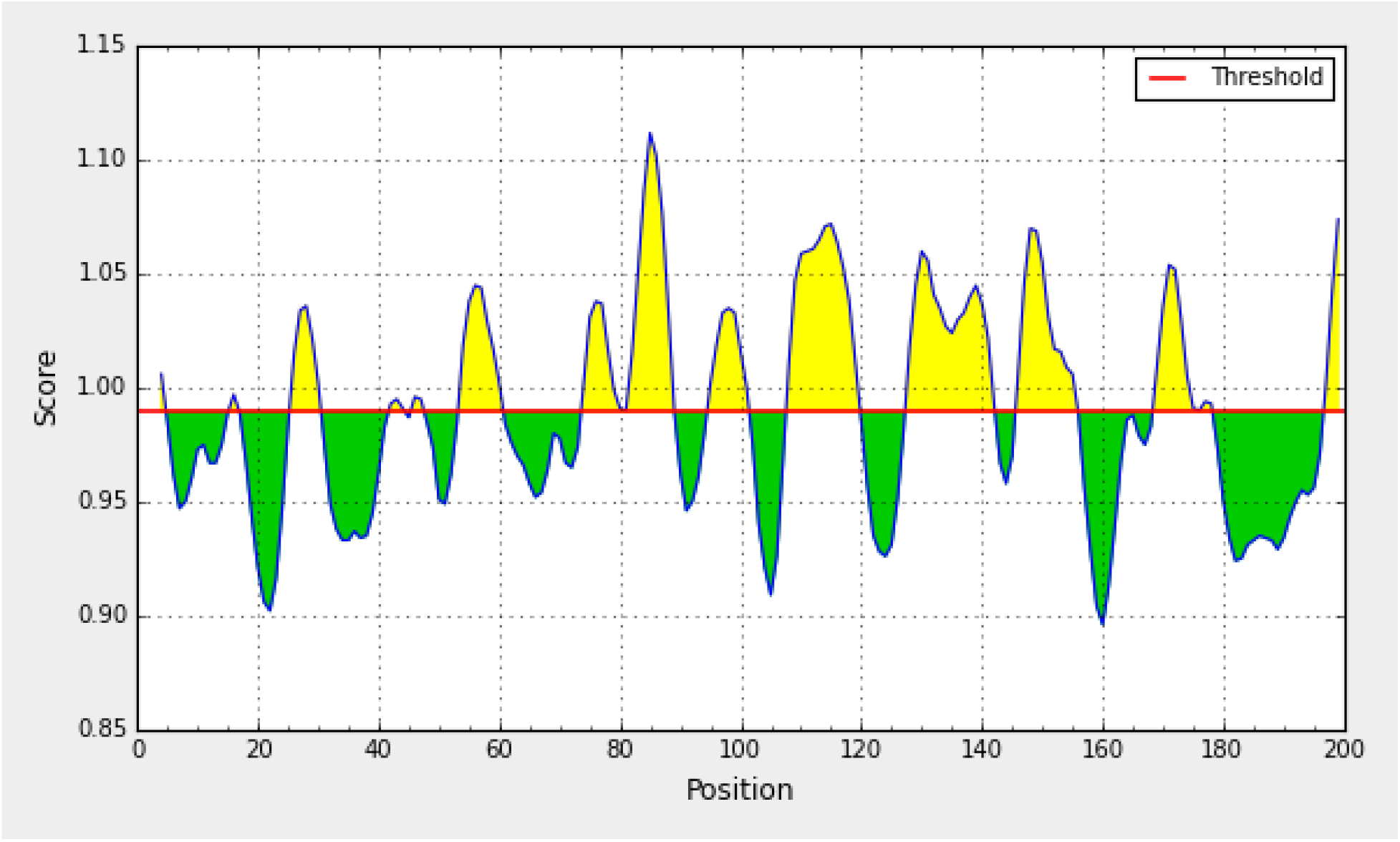
Karplus and Schulz flexibility prediction of a conserved region of spike glycoprotein of SARS-CoV-2.

#### 3.9.4. Prediction method of Bepipred linear epitope

This method uses hidden markov model (HMM) method (Best method for linear B-cell epitope prediction). The average antigenic propensity of the peptide was 0.075, with a maximum value of 1.896 and minimum value of 0.021. The threshold value of antigenic determination was 0.350. Peptide sequence from165 to 178 are capable of induction of the desired immune response from the B-cell epitope. The result is shown in Figure 8 and Table 8.

**Figure 8:**
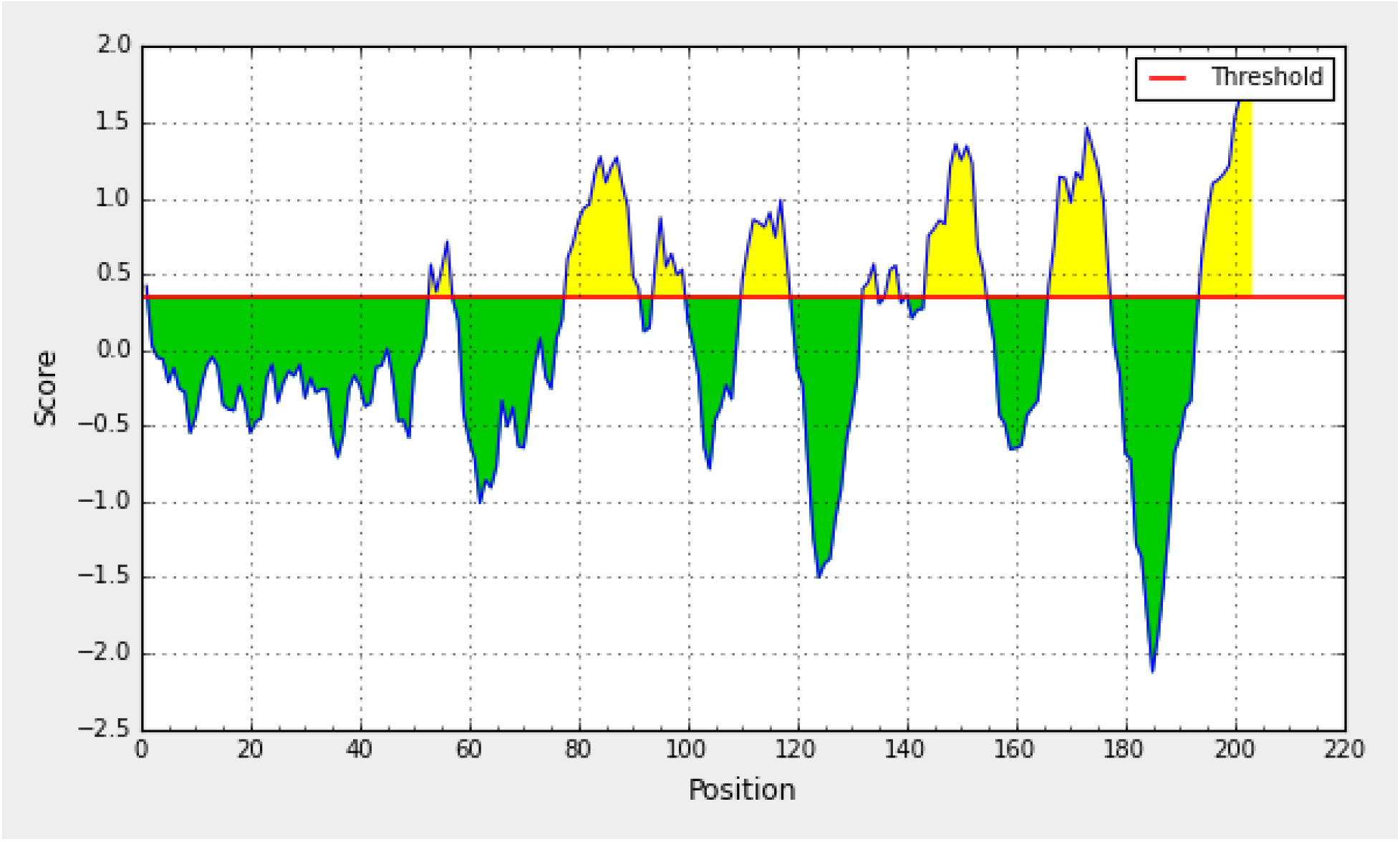
Bepipred linear epitope prediction of a conserved region of spike glycoprotein of SARS-CoV-2.

**Table 8:**
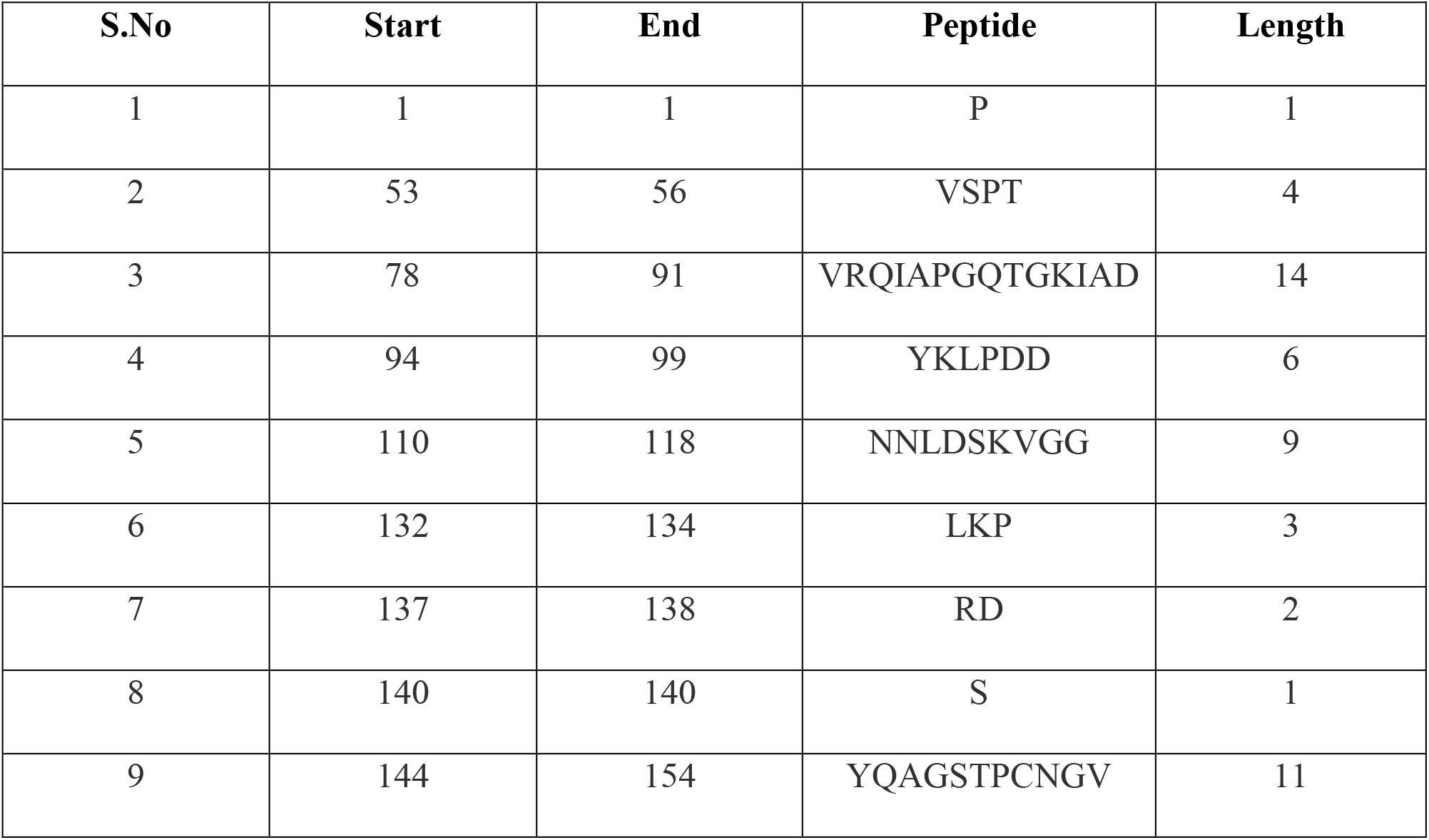
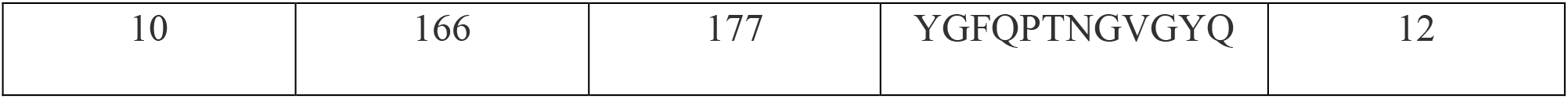
Shows B cell epitopes that are capable of producing a desired immune response, as predicted from Bepipred linear epitope analysis of the conserved region of spike glycoprotein of SARS-CoV-2.

#### 3.9.5. Prediction method of CF beta-turn

Often, the beta turns are hydrophilic and accessible. These 2 properties are of the antigenic region of a protein^7^ and so this method was used. The average value of antigenic propensity for peptide was 1.044, with a maximum value of 1.397 and minimum value of 0.694. The antigenic determination threshold was 1.044. The region of the peptide from 141 to 179 was considered as a beta turn region as shown in Figure 9.

**Figure 9:**
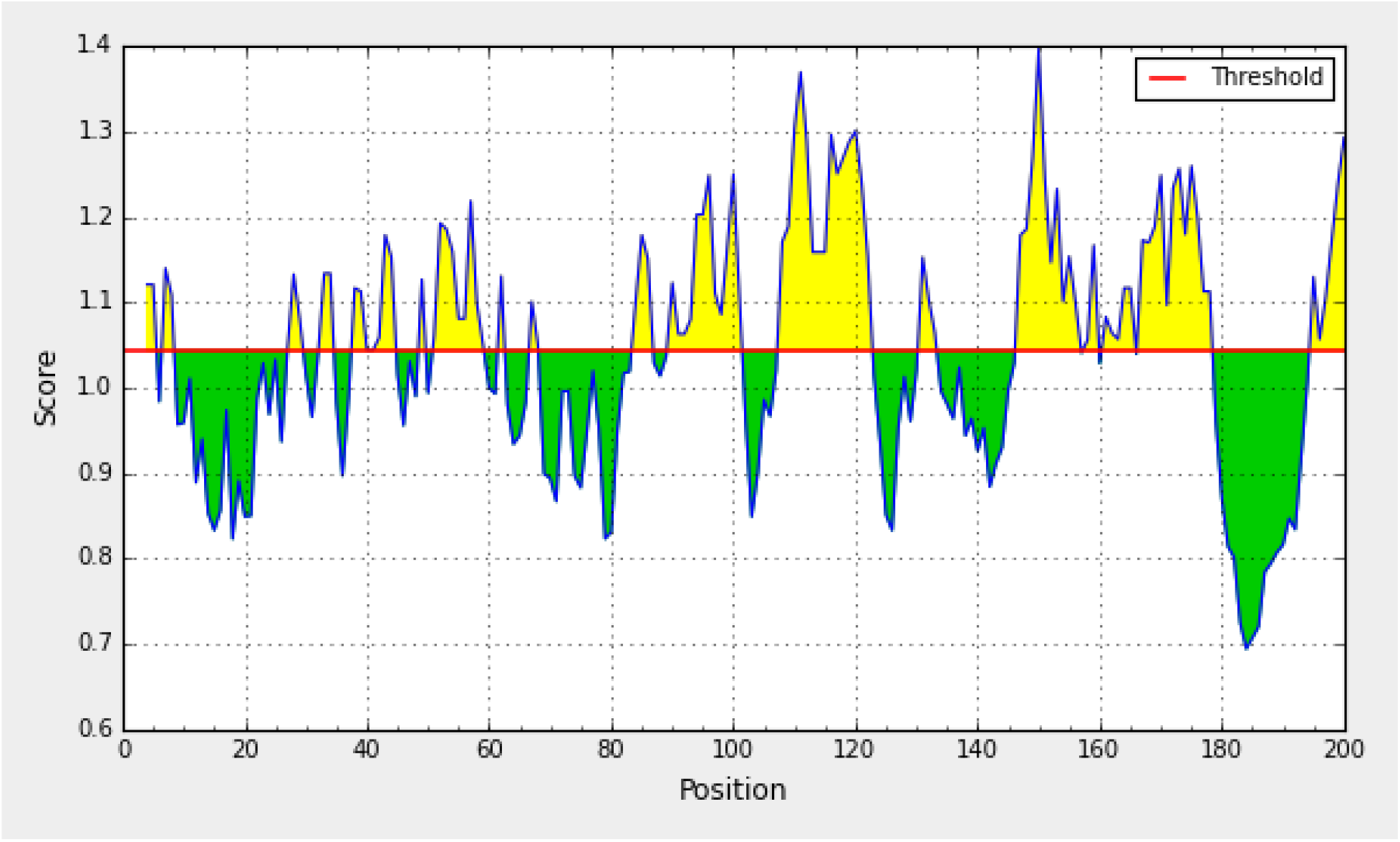
Chou and Fasman beta-turn prediction of a conserved region of spike glycoprotein of SARS-CoV-2.

## 4. DISCUSSION

Globally, SARS-CoV-2 infection has caused 728 013 deaths (as per the WHO report on August 10). This virus causes respiratory distress with other symptom like fever, dry cough and tiredness. Vaccine development plays an important role in elimination of the spreading virus^8–12^.

Historically, vaccination has saved lives of large human population from many pathogenic viruses. As compared to the traditional vaccination methods like live, killed and attenuated vaccines, epitope-based vaccine provide a much more rational method of vaccination. The reason behind this is that it contains a specific targeted immunogenic component of the pathogen that could elicit an immune response inside the host^13^. Examples of epitope-based vaccine design are available against rhinovirus^14^, dengue virus^15^, chikungunya virus^16^ and Saint Louis encephalitis virus^17^. As not much is known about the specific immunogenic response against SARS-CoV-2, recommendations of epitopes can be of great help in designing of epitope-based vaccines.

SARS-CoV-2 is a ribonucleic acid (RNA) virus that gets mutated more frequently^18^. These mutations mostly occur at the outer membrane S glycoprotein^19^ and increase the sustainability of the virus by escaping the humoral and cell-mediated immune response^20^. S protein can induce a faster and long-lived mucosal immune response, as compared to the other proteins of the virus^21^ and due to this fact, it is very popular among researchers^22, 23^. According to previous studies, S protein has proved to be an effective target for vaccine development against SARS-CoV and MERS-CoV virus^24, 25^. As virus RNA mutates, the conserved region of the S glycoprotein can be a very good target for the development of such an epitope-based vaccine against SARS-CoV-2 virus.

Mostly, vaccine developments are based on B-cell immunity. But, a strong immune response is generated by the CD8+ T-cell, as compared to the B-cell immunity^26^. Due to antigenic drift, with time antibody memory fails, whereas the T-cell immune response is long-lasting.

Due to the advancement in computational biology and sequence-based technology, there is a huge database that could be used in the treatment of such an infection. Therefore, an effort has been made in this paper to find a T-cell and B-cell epitope for designing of the epitope-based vaccine.

Although, the paper proposes a novel T-cell epitope but some B-cell epitopes are also being proposed. 23 T-cell epitopes with respective MHC-I molecules (having a value less than 200 of IC_50_) were found from immune database. These 23 T-cell epitopes were derived from the conserved S glycoprotein of SARS-CoV-2. Among them, 4 potential T-cell epitopes have fulfilled all the criteria that are important for a vaccine candidate epitope like stability, antigenicity, non-toxicity and non-allergenicity.

Further, good binding affinities have been found between the 4 selected epitopes and the MHC-I (having lowest values of IC_50_), whereas GQTGKIADY binds with HLA-C*03:03 most effectively. This means that GQTGKIADY could be the best possible epitope that could bind effectively with HLA-C*03:03, which is present inside the host system. Moreover, MD simulation results also support that the complex of GQTGKIADY and HLA-C*03:03 becomes stable after 40 ns.

Population coverage analysis was done and the results show that there is the presence of HLA-C*03:03 in the human population of India, China, Italy, United States, United Kingdom and Russia (countries that are mostly affected with SARS-CoV-2 infection). This means that GQTGKIADY epitope will bind effectively with HLA-C*03:03 that is present in the human population of the most effected countries with the infection of SARS-CoV-2, as mentioned above.

Further, many B-cell epitopes have also been found based on different properties and threshold that could elicit humoral response against SARS-CoV-2. These B-cell epitopes can also result in production of antibodies inside the host system. But we have focused mainly on the T-cell epitope for vaccine development, as T-cell epitope can elicit a long lasting immunity as compared to B-cell epitope.

This study is an in silico-based study and the data have been extracted from the various immune database, but such a type of study have previously been validated with wet-lab results^27^. So, the proposed B-cell and T-cell epitope could also be effective in eliciting an immune response and killing the infection caused by SARS-CoV-2.

## CONCLUSION

This study can help in designing of the epitope-based vaccine while saving a lot of wet lab effort. A novel T-cell epitope GQTGKIADY has been proposed as it contains all the features like antigenicity, non-allergenicity, non-toxicity and stability. Further, GQTGKIADY epitopes binds with HLA-C*03:03 very well, as predicted from the molecular docking and MD simulation results. Population coverage analysis results support that HLA-C*03:03 is present in the human population of the most effected countries like of India, China, Italy, United States, United Kingdom and Russia. This means the proposed T-cell epitope can be a very good candidate for eliciting the immune response in the most affected population. Some B-cell epitopes have also been found that could be effective in producing antibodies inside the host. This work mainly focuses on the T-cell epitopes, as T-cell based immune response could be long lasting as compared to B-cell for fighting with infection caused by SARS-CoV-2.

## ACKNOWLEDGEMENTS

We acknowledge the support of IIIT Allahabad for providing necessary facilities and infrastructure required for the completion of the work.

## References

1. Schoeman D, Fielding BC. Coronavirus envelope protein: current knowledge. Virol J 2019; 16(1):69.

2. Song W, Gui M, Wang X, Xiang Y. Cryo-EM structure of the SARS coronavirus spike glycoprotein in complex with its host cell receptor ACE2. PLoS Pathog 2018; 14(8):e1007236.

3. Remy V, Larger on N, Quilici S, Carroll S. The Economic Value of Vaccination: Why Prevention Is Wealth. Value Health J. Int. Soc. Pharmacoecon. Outcomes Res 2014; 17: A450.

4. Tenzer S, Peters B, Bulik S, Schoor O, Lemmel C, Schatz MM, et al. Modeling the MHC class I pathway by combining predictions of proteasomal cleavage, TAP transport and MHC class I binding. Cell Mol Life Sci 2005; 62:1025–1037.

5. Nair DT, Singh K, Siddiqui Z, Nayak BP, Rao KV, Salunke DM. Epitope recognition by diverse antibodies suggests conformational convergence in an antibody response. J Immunol 2002; 168(5):2371–2382.

6. Novotný J, Handschumacher M, Haber E, Bruccoleri RE, Carlson WB, Fanning DW, et al. Antigenic determinants in proteins coincide with surface regions accessible to large probes (antibody domains). Proc Natl Acad Sci USA 1986; 83(2):226–230.

7. Rose GD, Gierasch L, Smith JA. Turns in peptides and proteins. Adv Protein Chem 1985; 37:1–109.

8. Marshall SJ. Developing countries face double burden of disease. Bull. World. Health. Organ 2004; 82(7): 556.

9. De Groot, A.S.; Rappuoli, R. Genome-derived vaccines. Expert Rev Vaccines 2004; 3(1): 59–76.

10. Korber B, LaBute M, Yusim K. Immunoinformatics comes of age. PLoS Comput Biol 2006; 2(6): e71.

11. Fauci AS. Emerging and re-emerging infectious diseases: influenza as a prototype of the host-pathogen balancing act. Cell 2006; 124(4): 665–670.

12. Purcell AW, McCluskey J, Rossjohn J. More than one reason to rethink the use of peptides in vaccine design. Nat Rev Drug Discov 2007; 6(5): 404–414.

13. Patra P, Mondal N, Patra BC, Bhattacharya M. Epitope-Based Vaccine Designing of Nocardia asteroides Targeting the Virulence Factor Mce-Family Protein by Immunoinformatics Approach. Int J Pept Res Ther 2019; 26(2): 1165–1176.

14. Lapelosa M, Gallicchio E, Arnold GF, Arnold E, Levy RM. In silico vaccine design based on molecular simulations of rhinovirus chimaeras presenting HIV-1 gp41 epitopes. J Mol Biol 2009; 385(2):675–691.

15. Chakraborty S, Chakravorty R, Ahmed M, Atiqur R, Zaved WT, Faizule H, et al. A computational approach for identification of epitopes in dengue virus envelope protein: a step towards designing a universal dengue vaccine targeting endemic regions. In Silico Biol 2010; 10(5–6):235–246.

16. Islam R, Sakib MS, Zaman A. A computational assay to design an epitope-based peptide vaccine against chikungunya virus. Future Virol 2012; 7(10):1029–1042.

17. Hasan MA, Hossain M, Alam MJ. A computational assay to design an epitope-based Peptide vaccine against Saint Louis encephalitis virus. Bioinform Biol Insights 2013; 7:347–355.

18. Twiddy SS, Holmes EC, Rambaut A. Inferring the rate and timescale of dengue virus evolution. Mol Biol Evol 2003; 20(1):122–129.

19. Manzin A, Solforosi L, Petrelli E, Macarri G, Tosone G, Piazza M, et al. Evolution of hypervariable region 1 of hepatitis C virus in primary infection. J Virol 1998; 72(7): 6271–6276.

20. Christie JM, Chapel H, Chapman RW, Rosenberg WM. Immune selection and genetic sequence variation in core and envelope regions of hepatitis C virus. Hepatology 1999; 30(4):1037–1044.

21. Ma C, Li Y, Wang L, Zhao G, Tao X, Tseng CK, et al. Intranasal vaccination with recombinant receptor-binding domain of MERS-CoV spike protein induces much stronger local mucosal immune responses than subcutaneous immunization: Implication for designing novel mucosal MERS vaccines. Vaccine 2014; 32(18):2100–2108.

22. Yang ZY, Kong WP, Huang Y, Roberts A, Murphy BR, Subbarao K, et al. A DNA vaccine induces SARS coronavirus neutralization and protective immunity in mice. Nature 2004; 428(6982):561–564.

23. Agnihothram S, Gopal R, Yount BL Jr, Donaldson EF, Menachery VD, Graham RL, et al. Evaluation of serologic and antigenic relationships between middle eastern respiratory syndrome coronavirus and other coronaviruses to develop vaccine platforms for the rapid response to emerging coronaviruses. J Infect Dis 2014; 209(7):995–1006.

24. Du L, He Y, Zhou Y, Liu S, Zheng BJ, Jiang S. The spike protein of 353 SARS-CoV--a target for vaccine and therapeutic development. Nat Rev Microbiol 2009; 7:226–236.

25. Schindewolf C, Menachery VD. Middle East Respiratory Syndrome Vaccine Candidates: Cautious Optimism. Viruses 2019; 11(1):74.

26. Shrestha B, Diamond MS. Role of CD8+ T cells in control of West Nile virus infection. J Virol 2004; 78(15):8312–8321.

27. Khan MK, Zaman S, Chakraborty S, Chakravorty R, Alam MM Bhuiyan TR, et al. In silico predicted mycobacterial epitope elicits in vitro T-cell responses. Mol Immunol 2014; 61(1):16–22.

